# A Photoactivated Protein Degrader for Optical Control of Synaptic Function

**DOI:** 10.1101/2023.02.13.528397

**Authors:** T. Ko, C. Jou, A.B. Grau-Perales, M. Reynders, A.A. Fenton, D. Trauner

**Affiliations:** Department of Chemistry, University of Pennsylvania, 231 South 34th Street Philadelphia, PA 19104-6323, USA; Department of Psychology, Hunter College, 695 Park Avenue, New York, NY, 10065, USA; Center for Neural Science, New York University, 4 Washington Place, New York, NY 10003, USA; Department of Chemistry, New York University, 100 Washington Square East, New York, NY 10003, USA

## Abstract

Hundreds of proteins determine the function of synapses, and synapses define the neuronal circuits that subserve myriad brain, cognitive, and behavioral functions. It is thus necessary to precisely manipulate specific proteins at specific sub-cellular locations and times to elucidate the roles of particular proteins and synapses in brain function. We developed PHOtochemically TArgeting Chimeras (PHOTACs) as a strategy to optically degrade specific proteins with high spatial and temporal precision. PHOTACs are small molecules that, upon wavelength-selective illumination, catalyze ubiquitylation and degradation of target proteins through endogenous proteasomes. Here we describe the design and chemical properties of a PHOTAC that targets Ca^2+^/calmodulin-dependent protein kinase II alpha (CaMKIIα), which is abundant and crucial for baseline synaptic function of excitatory neurons. We validate the PHOTAC strategy, showing that the **CaMKIIα-PHOTAC** is effective in mouse brain tissue. Light activation of **CaMKIIα-PHOTAC** removed CaMKIIα from regions of the mouse hippocampus only within 25 μm of the illuminated brain surface. The optically-controlled degradation decreases synaptic function within minutes of light activation, measured by the light-initiated attenuation of evoked field excitatory postsynaptic potential (fEPSP) responses to physiological stimulation. The PHOTACs methodology should be broadly applicable to other key proteins implicated in synaptic function, especially for evaluating their precise roles in the maintenance of long-term potentiation and memory within subcellular dendritic domains.

## Introduction

The ability to perturb individual components of complex networks with light has enabled unprecedented progress in neuroscience. Optical excitation through light-gated ion channels like channelrhodopsin and light-gated chloride pumps like halorhodopsin, is now widely employed for untangling the diverse roles of individual neurons and defining functionally-distinct neuronal circuits defined by effective synapses ^1^. Channelrhodopsin, halorhodopsin and other optogenetic effector molecules are powerful research tools because they are genetically encoded, allowing the precisely timed manipulation of neurons and other cell types that express them. However, certain applications and particular brain functions, for instance, the maintenance of long-term memory in hippocampus, depend on subcomponents of neurons, such as a synaptic compartment or subsets of synapses. Although such subcellular specificity cannot be sufficiently targeted by genetic identity, the specificity can be achieved in combination with precise delivery of light ^2-4^.

Some of the most intensely studied processes in neuroscience are ‘when,’ ‘where,’ and ‘how’ memories are formed, consolidated, retrieved, and their storage maintained. While it is not universally agreed that protein-mediated synaptic plasticity is the neurobiological basis of memory ^5^, there is compelling evidence that certain proteins play crucial, temporally-restricted roles in both activity-dependent long-term potentiation (LTP) of synaptic function and also memory. Both are dynamic processes with mechanistically distinct phases ^6-8^.

Amongst many memory-shaping proteins, Ca^2+^/calmodulin-dependent protein kinase II alpha (CaMKIIα) is of particular interest. It is ubiquitously expressed in excitatory neurons throughout the mammalian forebrain, including the hippocampus. Once activated by calcium influx, for example upon repeated postsynaptic stimulation of NMDA receptors, the dodecameric CaMKIIα stays activated through continuous autophosphorylation. CaMKIIα is crucial for maintaining basal synaptic function, and as a consequence, also for maintaining synaptic function after LTP-inducing stimulation and long-term memory training ^9-11^. CaMKIIα is necessary for the protein synthesis-independent induction of LTP (early-LTP) and learning, the acquisition of memory; CaMKIIα inhibition may no longer impair either process once protein-synthesis dependent late-LTP and/or long-term memory has been established, which is why we targeted CaMKIIα and baseline synaptic function to test and validate a novel methodology for manipulating synaptic function by a photoactivated degrader of proteins crucial for synaptic function.

Here, we report a method to precisely control the level of a protein using light, which we validate by targeting CaMKIIα. Our method is not based on manipulating genetic expression, such as gene knockout technologies or knockdown by RNA manipulations, nor is our approach based on occupancy driven pharmacology, each of which have limitations for studying persistent memory kinases. This is due to confounds that include genetic compensation, crude temporal and spatial control, and concentration-dependent off-target effects ^12-14^. Neither does our approach involve an engineered protein, the over expression of which might lead to unexpected and undesired physiological responses ^15-17^.

Our strategy rather relies on harnessing the cell’s protein degradation machinery with PROteolysis TArgeting Chimeras (PROTACs). These are bifunctional molecules that combine a ligand for an E3 ubiquitin ligase with a second ligand that targets a protein of interest (POI), in this case CaMKIIα. The ligands promote tertiary complex formation, the polyubiquitination of the POI, and its subsequent proteasomal degradation ^18^. We have endowed PROTACs with a molecular photoswitch that makes them inactive in the dark but makes them active degraders of the POI upon illumination with deep violet light that has limited tissue penetration. The resulting systems are called PHOTACs (PHOtochemically TArgeting Chimeras), and in cell culture have already been demonstrated to degrade BET proteins (BRD2-4), and FKBP12 in a light dependent fashion ^19^. They are based on a photoswitchable ligand for the E3 ligase cereblon (CRBN) that is widely expressed in the brain ^20^ and they can be easily modified with respect to the POI. We now show that the PHOTACs strategy can be extended to brain tissue and to CaMKIIα, a persistent kinase, crucial for synaptic function.

## Results

### Design and characterization of a CaMKIIα-PHOTAC

The design of our PHOTAC was based on ligands that have been structurally characterized in conjunction with CaMKIIα in its open and closed form ^21^. These include bosutinib and indirubin derivative E804, which bind to the kinase domain and inhibit it, and 5-hydroxy diclofenac (5-HDC), which binds to the γ-hydroxybutyrate site in the CaMKIIα hub domain and does not inhibit kinase activity ^22-24^. We were particularly interested in bosutinib as inspection of its X-ray structure bound to CaMKIIα reveals a solvent exposed piperazine motif that would be a suitable site for ligand attachment without disrupting crucial binding interactions ^21, 25^ (Fig. 1B). This prompted us to prepare a bosutinib-based PHOTAC with the primary aim of degrading CaMKIIα. The structure of this PHOTAC, which we term **CaMKIIα-PHOTAC**, and its photoswitching parameters are shown in Fig. 1. Its synthesis is shown in Fig. S1A. A control compound, **Me-CaMKIIα-PHOTAC**, wherein the imide moiety is *N*-methylated, was also synthesized (Fig. S1B). *N*-Methyl imides are unable to form ternary complexes with CRBN despite a minor modification in molecular composition ^26^.

**Figure 1.**
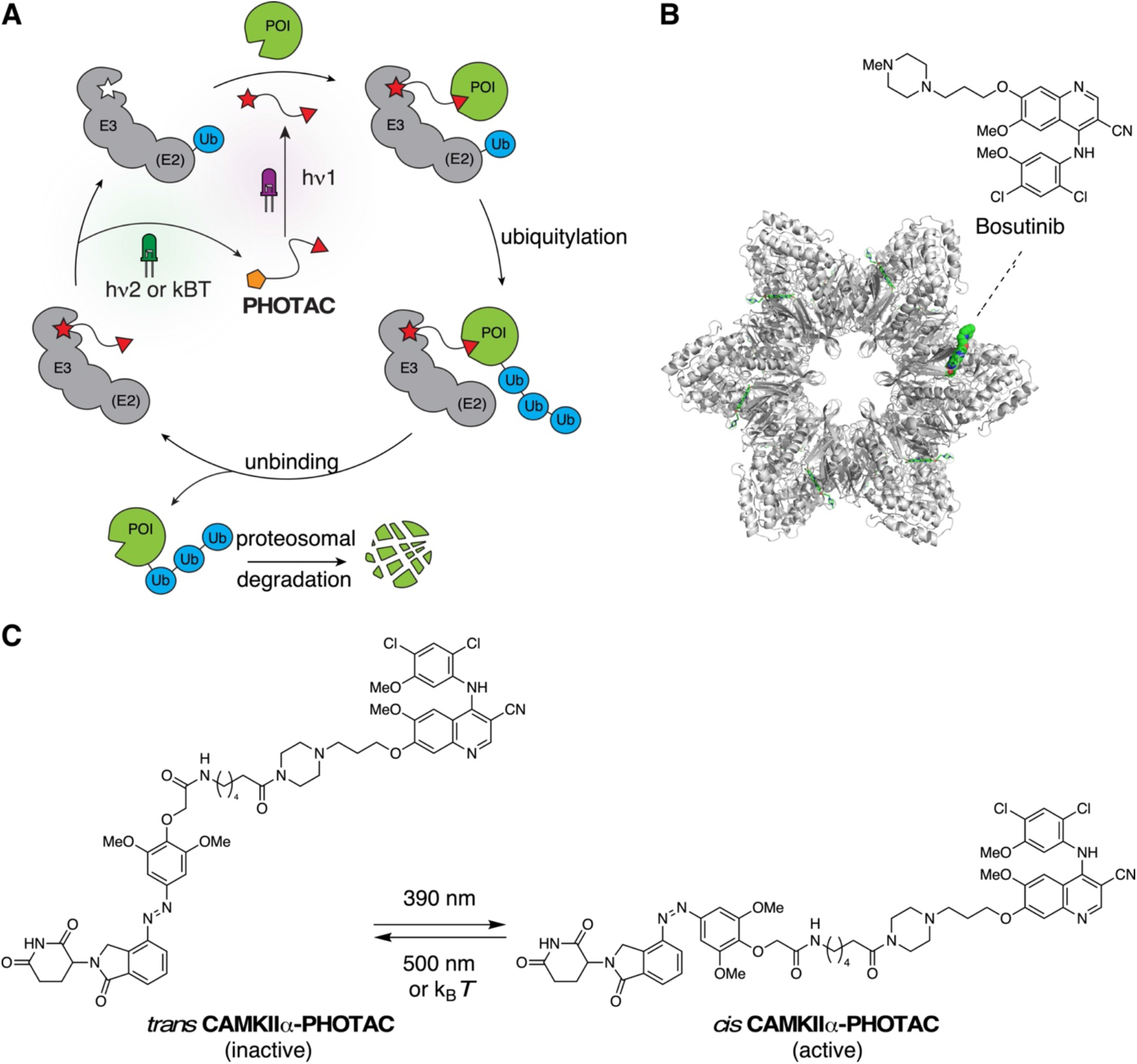
Logic of PHOTACs and molecular design of a CaMKIIα-PHOTAC. A) Schematic depiction of a PHOTAC. The E3 ligase ligand can be toggled between an inactive (orange pentagon) and an active (red star) form upon irradiation. Activation of PHOTAC leads to ternary complex formation with both an E3 ligase and a protein of interest (POI), and the subsequent degradation of the POI. B) Dodecametric CaMKIIα holoenzyme crystal structure bound to bosutinib (PDB: 3SOA). C) The CaMKIIα-PHOTAC that can be switched between its thermodynamically more stable and inactive *trans* isomer (left) and active *cis* isomer (right).

### Effects of light activation of CaMKIIα-PHOTAC on CaMKIIα

We then investigated the effect of **CaMKIIα-PHOTAC** on CaMKIIα using Western Blot analysis (Fig. S2). Dorsal hippocampus slices were cut 300 μm thick, then incubated for 6 hours in a artificial cerebrospinal fluid (aCSF) recovery buffer with 3 μM **CaMKIIα-PHOTAC** with or without 390-nm 100-ms illumination pulses every 10 seconds for 30 minutes ^27^. The tissue was lysed and homogenized, and the resulting protein lysate was separated through gel electrophoresis and blotted against CaMKIIα, which is an abundant protein accounting for 2% of all hippocampal proteins ^28^. CaMKIIα levels were indistinguishable with or without light activation (F_2,14_ = 0.67, p = 0.53). Because the tissue penetration of 390 nm light is limited compared to longer wavelengths ^29^, this lack of effect in Western blots indicated that the **CaMKIIα-PHOTAC** was either ineffective, or alternatively, the degradation of abundant CaMKIIα was tightly limited spatially.

We used immunohistochemistry and optical sectioning confocal microscopy to distinguish the two possibilities. Dorsal hippocampal slices were prepared and treated in the same way with 3 μM **CaMKIIα-PHOTAC** with or without 45 min of 390 nm pulsed illumination; control slices also received the 390 nm illumination. The tissue was immediately fixed, prepared for DAPI staining to assess cellular integrity, and immunohistochemical staining against CaMKIIα and both MAP2 and parvalbumin as control proteins was carried out. CaMKIIα immunostaining was significantly decreased in the 390 nm illuminated samples treated with the CaMKIIα-PHOTAC (n=10), compared to the unilluminated (n=8) and the aCSF + 390 nm (n=7) and aCSF + dark (n=4) control slices (Fig. 3A-C); 1-way ANOVA F_3,25_ = 9.0, p = 0.0002, aCSF+ dark = aCSF+ 390 nm = PHOTAC + dark > PHOTAC + 390 nm). In contrast, no significant group differences in either DAPI (F_3,25_ = 0.053, p = 0.98), MAP2 (F_3,13_ = 0.064, p = 0.98) or parvalbumin (F_3,11_ = 0.036, p = 0.99) signals were detected, indicating no evidence of cellular toxicity or nonspecific effects of activating the **CaMKIIα-PHOTAC**. The control compound **Me-CaMKIIα-PHOTAC** also had no effect on CaMKIIα immunostaining (Fig. S3). In combination with observing no effects on western blotting despite using the same primary antibody, these findings indicate that light-activated **CaMKIIα-PHOTAC** is effective in selectively destroying CaMKIIα in a spatially-limited domain.

Using confocal optical sectioning, we then evaluated the spatial extent of CaMKIIα loss by the light-activated CaMKII-PHOTAC along the direction of incident light (Fig. 4A). Dorsal hippocampus slices were prepared, treated with CaMKIIα-PHOTAC or aCSF and then 390-nm light or dark illuminated, as before, followed by immunohistochemistry (PHOTAC + 390 nm n = 7, PHOTAC + dark n = 4, aCSF + 390 nm n = 5, aCSF + dark n = 4). Confocal microscopy identified CaMKIIα immunostaining at 0, -12.5, and -25 μm depths relative to the surface of the slice. CaMKIIα levels were unchanged across the examined depths in control treated slices, whether or not they were illuminated with 390 nm light as well as the **CaMKIIα-PHOTAC** treated slices that were not illuminated. In contrast, CaMKIIα immunostaining was weakest at the surface (0 μm), intermediate at 12.5 μm and not significantly changed at the 25 μm depth, although there was a trend to be reduced (Fig. 4B,C). Analysis of the raw cell count values confirmed these impressions. The effects of the treatment (F_3,16_ = 9.71, p = 0.0007), depth (F_2,16_ = 43.39, p = 10^−7^), and the treatment X depth interaction (F_6,16_ = 11.34, p = 10^−5^) were all significant. The CaMKIIα loss in the PHOTAC + 390 nm slices appeared to decrease linearly with depth, motivating us to fit the CaMKIIα-positive cell counts at the three depths of each slice by linear regression. Accordingly, we calculated the percent change relative to the average CaMKIIα level at 25 μm in the control slices (Fig. 4C) and linearly fit the three relative measurements (r’s = 0.52 - 0.99, p’s < 0.05). The regression slopes describe the CaMKIIα loss as a function of depth and the 1-way ANOVA comparing the slopes of the four conditions confirmed significant effects of condition (F_3,16_ = 13.13, p = 0.0001, aCSF + dark = aCSF + 390 nm = PHOTAC + dark > PHOTAC + 390 nm). The average slope (−1.87±0.25 %/μm) for the PHOTAC + 390 nm condition indicates that at the illumination intensity we use, the effect on CaMKIIα is limited to within 50 μm. Accordingly, the CaMKIIα loss should be less than 7% of the 300 μm hippocampus slices, which can explain why the loss of CamKIIα was not detected by western blot. We conclude that the CaMKIIα-PHOTAC is effective within a limited spatial domain.

### Effects of light activation of CaMKIIα-PHOTAC on synaptic function

We used electrophysiology to test whether there are functional effects of spatially-limited degradation of CaMKIIα following light activation of the **CaMKIIα-PHOTAC**. Because CaMKIIα is crucial for basal excitatory synaptic function^9^, the predicted effect of activating **CaMKIIα-PHOTAC** is decreased synaptic responses to test electrophysiological stimulation. Furthermore, by continuously monitoring synaptic responses to test stimulation we could determine the time course of the PHOTAC effect. Dorsal hippocampus slices were prepared as in the biochemistry and immunohistochemistry experiments and incubated 6 hours in an artificial cerebral spinal fluid (aCSF) recovery buffer with or without 3 μM CaMKIIα-PHOTAC. All slices were moved to a submerged chamber for electrophysiological recording of baseline synaptic responses to a 100-μs, ∼200 mA stimulus pulse that elicited a 70% maximum response (Fig. 5B). The test stimulus was delivered every 30 s to the perforant path, the evoked field excitatory postsynaptic potential (fEPSP) response in the supra-pyramidal blade of the dentate gyrus was recorded, and its slope was measured and normalized to each slice’s average baseline response. After recording 10 minutes of basal responses, at time = 0 s, a train of 385 nm 100-ms light pulses every 10 s was initiated in the **CaMKIIα-PHOTAC** treated and the aCSF control slices. A separate set of **CaMKIIα-PHOTAC** treated slices did not receive the light stimulation (PHOTAC+Dark). By inspection, single slice experiments show that while neither the **CaMKII**α**-PHOTAC** treatment itself (PHOTAC+Dark), nor the light illumination itself (aCSF+385 nm) changes synaptic responses, the illumination depresses baseline synaptic responses in the PHOTAC-incubated slices (PHOTAC+385 nm), and the effect is detected in under five minutes, as predicted by light-activated **CaMKIIα-PHOTAC** mediated destruction of CaMKIIα (Fig. 5B). The group comparison confirms that a significant decrease in synaptic strength was only observed when slices received both the **CaMKIIα-PHOTAC** and illumination. In contrast, there was no overall change in the absence of either the PHOTAC or illumination (Fig. 5C). The average normalized fEPSP responses were analyzed in each 5-min epoch (10 responses) (Fig. 5C). Responses were stable and equivalent during the two 5-min baseline epochs (Treatment: F_2,18.6_ = 0.03, p = 0.97; Time: F_1,18.6_ = 1.40, p = 0.25; interaction: F_2,18.6_=0.45, p = 0.64). Light stimulation during 45 minutes to activate the PHOTAC was effective only in the PHOTAC+385 nm Treatment (Fig. 5D; F_2,75.6_ = 55.14, p = 10^−16^; PHOTAC+385 nm < PHOTAC+Dark = aCSF+385 nm), with no significant effects of time (F_9,41.4_ = 0.31, p = 0.97) or the interaction (F_18,52.1_ = 1.68, p = 0.07).

## Discussion

### Summary, Features and Limitations of the CaMKIIα-PHOTAC

We performed a proof-of-concept study, demonstrating for the first time the feasibility of a reversible PHOTAC for optical control of selective protein degradation in neural tissue (Figs. 1-3). PHOTACs have previously only been demonstrated to be effective in cell culture ^19^. The **CaMKIIα-PHOTAC** was designed to recruit the native E3 ligase cereblon in response to short wavelengths, which limited tissue penetration along the direction of incident light ^19^. The effect of the PHOTAC was detectable by IHC to an imaging depth of 12.5 μm, but not 25 μm (Fig. 4). Assaying synaptic function allowed us to confirm that the PHOTAC is ineffective until activated by the proper wavelength of light, after which protein degradation is rapid; the physiological effects could be detected within five minutes, consistent with the design expectations of the PHOTACs strategy to harness the native ongoing proteasomal mechanism of protein degradation (Fig. 5). The light-activated decline in synaptic function was not accompanied by evidence of cell loss or unintended damage as DAPI staining for nuclei, and measures of cytoskeletal protein MAP2, and calcium binding protein parvalbumin were unaffected in the same slices that expressed a 3-fold decrease of CaMKIIα (Fig. 3). Crucially we demonstrate that the strategy was able to restrict the loss of CaMKIIα to within 25 μm of the illumination (Fig. 4,), in contrast to genetic knockout and knockdown strategies that operate throughout a cell. Recent investigations into the contested relationship between changes in synaptic strength and storage of information in memory ^5^ have identified that that memory-associated synaptic change can be restricted to particular dendritic compartments ^30^, and at particular synapses within these compartments ^31^, making as we have demonstrated, sub-cellular manipulations of the hypothesized protein components of memory storage essential for progress. We conclude the **CaMKIIα-PHOTAC** was both selective and effective, demonstrating feasibility of the PHOTACs strategy for selective protein degradation at subcellular spatial resolution.

### Other protein targets

The PHOTACs strategy can target any protein of interest so long as there is a selective ligand, even if the ligand is a low affinity inhibitor of protein function because the PHOTAC mechanism is to catalyze ubiquitination of the protein of interest. In principle, a single PHOTAC molecule can catalyze the ubiquitination and subsequent destruction of many thousands of proteins. Consequently, the PHOTACs strategy should be effective at concentrations that are substantially lower than the effective concentration for functional stoichiometric inhibition, even if the PHOTAC ligand also functions as an inhibitor. Indeed, at a 3 μM concentration, we observed a lack of any molecular, cellular, or physiological effect of the control **Me-CaMKIIα-PHOTAC** and **CaMKII - PHOTAC** prior to light activation, despite both molecules being comprised of bosutinib, which is a tyrosine kinase inhibitor with effects beyond CaMKIIα on PI3K/AKT/mTOR, MAPK/ERK, and JAK/STAT3 signaling.^32.^ In hippocampal neurons, CaMKII*α* is much more abundant than the kinases in these pathways. A functional PROTAC was reported utilizing bosutinib as a ligand to induce degradation of c-ABL and the oncogenic fusion protein BCR-ABL ^33^, which have roles in neuroinflammation ^34^ and development of neuronal processes through microtubule associated protein (MAP) kinase signaling ^35^. Although degradation of native c-ABL expressed in brain tissue could be expected by activating the **CaMKIIα-PHOTAC**, we observed no changes in MAP2 expression, and thus no evidence of effects on c-ABL (Fig 3F). While we cannot rule out extra-CaMKIIα effects, none manifested in our assays designed to assess effects on distinct levels of biological organization.

### Potential application-specific modifications

We have described the concept of a general-purpose strategy to degrade proteins of interest under optical control and demonstrated feasibility. Going forward, PHOTACs will be designed to meet particular specifications as dictated by a particular experimental question. For example, the characteristics of the **CaMKIIα-PHOTAC** we designed are advantageous for experiments that call for limiting where the target protein is degraded upon light activation, but not for widespread protein degradation. An example experiment might be to test whether turnover and/or upregulation of a dendritic synaptic protein is due to local translation, rather than transport from distant sites. Indeed, the use of antisense oligonucleotides previously allowed us to determine that both LTP and learning increase newly synthesized protein kinase M zeta (PKMζ). While we hypothesize the PKMζ increases would result from local translation, it has not been demonstrated ^13, 36^. This could be determined by using short wavelength light to activate an appropriately designed PKMζ-PHOTAC selectively at or between the soma and the synaptic input compartment of a hippocampal slice. In contrast, to determine if loss of PKMζ erases memory (or established LTP *in vivo*), an effectively designed PKMζ-PHOTAC would photoactivate in response to more tissue penetrant, longer wavelengths. Genetically encoded PHOTACs are possible, and these will be the focus of our upcoming activities. Indeed, ubiquitously expressing a PHOTAC with spatially limited activation properties like we have demonstrated, or with cell-type specificity can in principle, be powerfully combined with spatially precise two-photon (2P) excitation of spatial targets like a dendritic compartment or even specific dendritic subcomponents, using light sculpting and holographic illumination techniques ^37^. Effective 2P activation of azobenzene photoswitches has been demonstrated ^38^. Spatial precision can be further enhanced by taking advantage of the demonstrated PHOTAC reversibility implemented with spectrally-specific activation and inactivation of protein destruction (Fig. 2D) organized as center-surround or similar contrast-enhancing antagonistic patterns.

**Figure 2.**
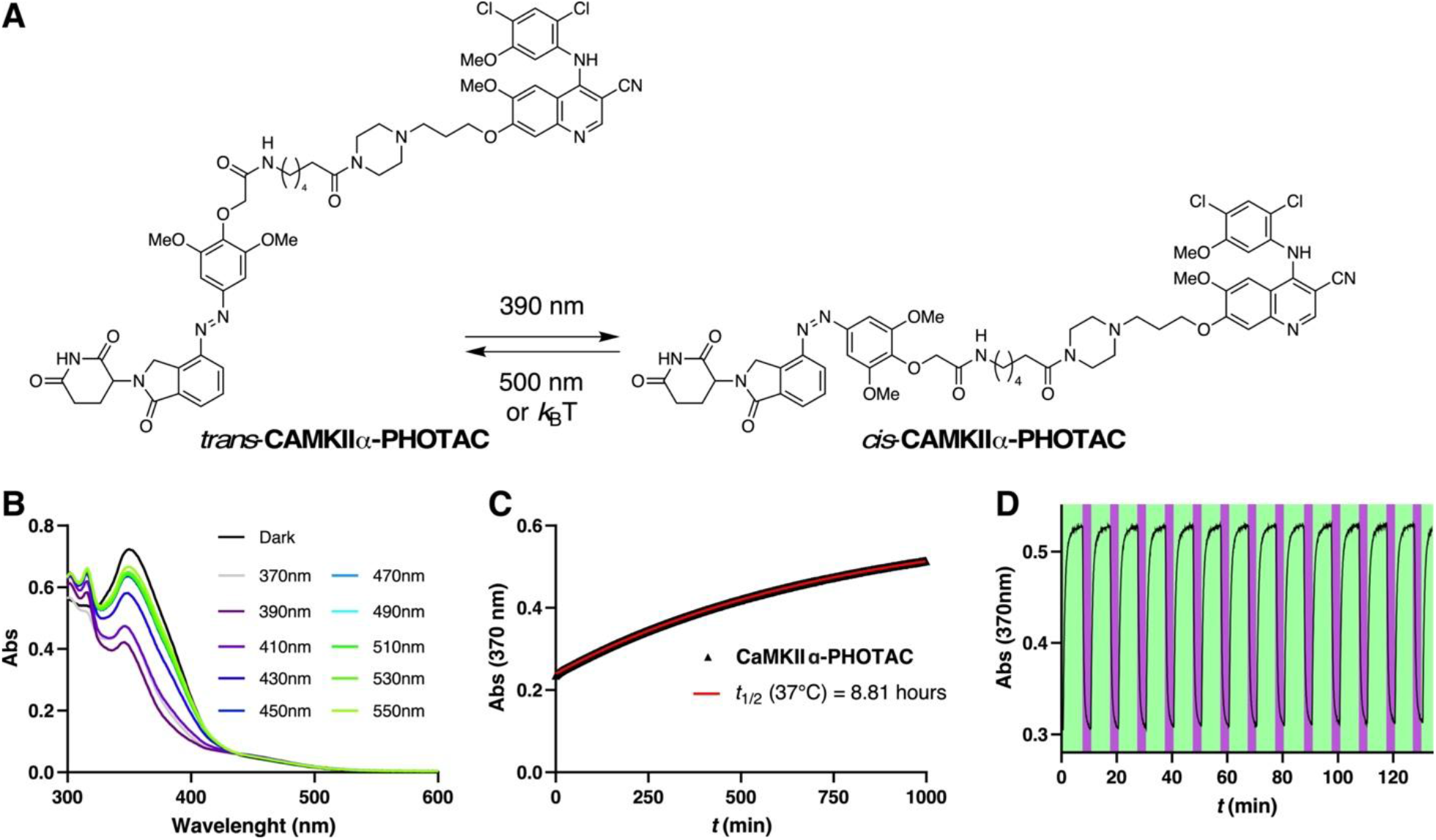
Photophysical characterization of CaMKIIα-PHOTAC. A) Isomerization of *trans*-**CaMKIIα-PHOTAC** to *cis*-**CaMKIIα-PHOTAC**. B) UV-Vis spectra of CaMKIIα-PHOTAC (50 μM) in the dark and at different photostationary states in DMSO at r.t. C) Thermal relaxation of CaMKIIα-PHOTAC (50 μM) at 37 °C in DMSO D) Reversible *trans* → *cis* isomerization of CaMKIIα-PHOTAC (50 μM) at 390 (purple):500 (green) nm irradiation in DMSO at r.t.

**Figure 3.**
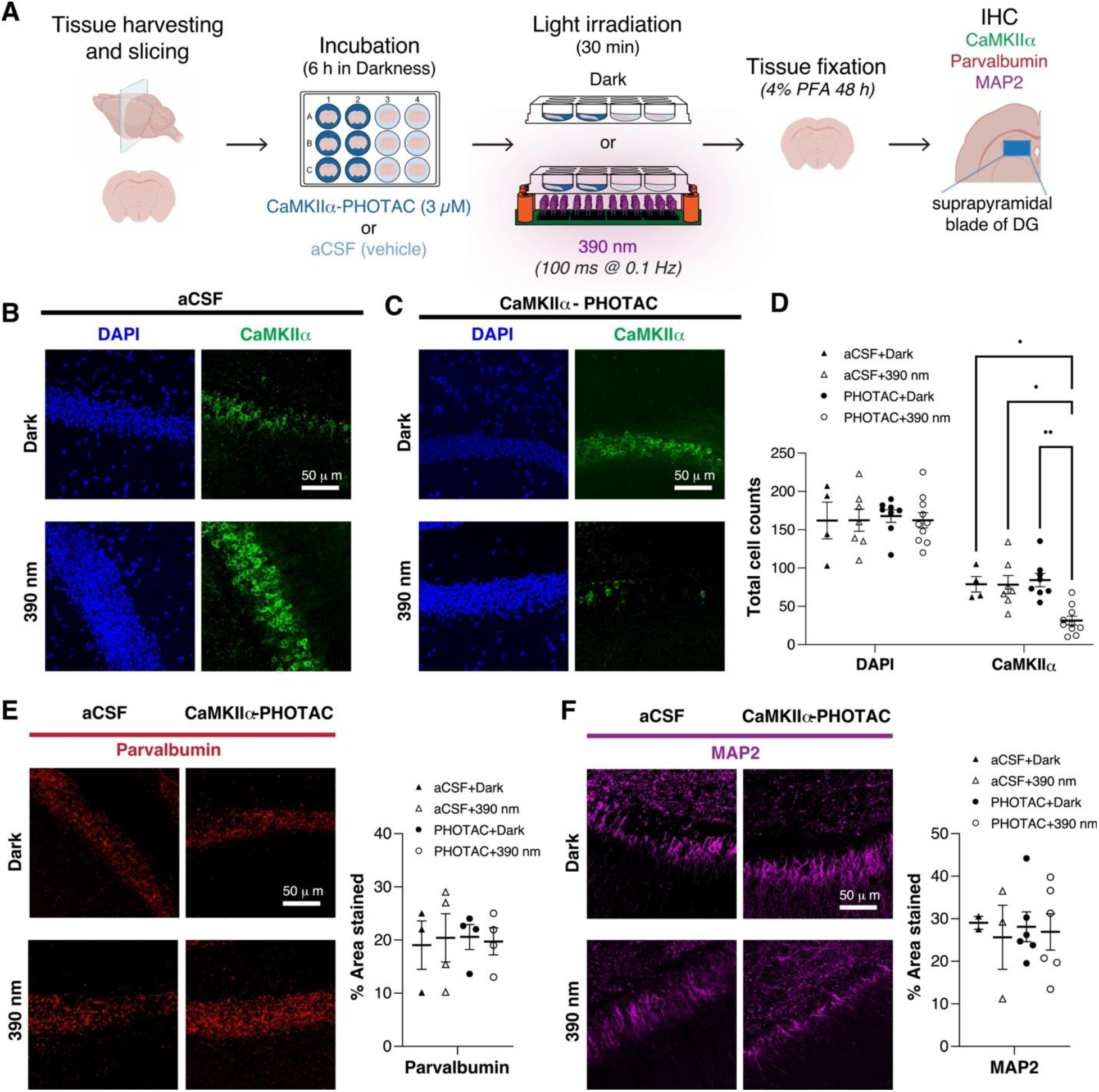
CaMKIIα degradation in brain slices monitored by immunohistochemistry. A) Assay schematic. Mouse brain slices were harvested and incubated with either CaMKIIα-PHOTAC (3 μM) or aCSF (vehicle). After light irradiation (390 nm, 100 ms every 10 s) or incubation in the dark, the tissue was fixed and immunostained. B-D) Examples and quantitation of DAPI staining and immunohistochemistry for CaMKIIα. E) Parvalbumin, and F) MAP2. *p < 0.05, **p< 0.01 post-hoc tests.

**Figure 4.**
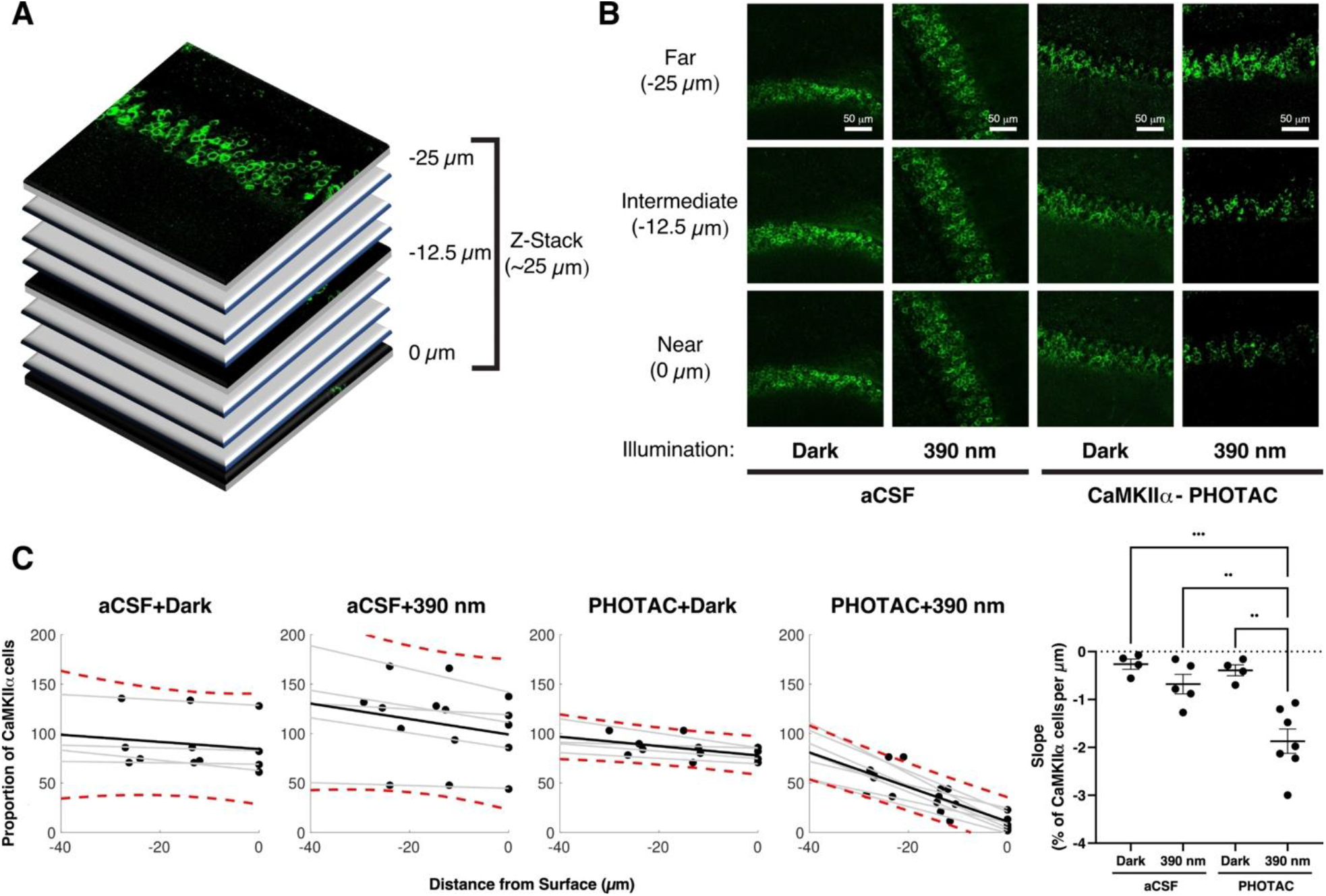
The CaMKIIα-PHOTAC mediated loss of CaMKIIα is spatially limited along the direction of incident light. A) Confocal optical sections were evaluated from the surface of 300-μm hippocampal slices that were prepared for CaMKIIα-immunohistochemistry using 25-μm-depth Z-Stacks on an upright Leica SP8 confocal microscope. B) From the 25-μm-depth Z-Stacks, the amount of CaMKIIα was measured at the most superficial layer, the middle layer, and the bottom layer (Near, Intermediate and Far, respectively) of each Z-Stack and C) quantitation of CaMKIIα in optical sections from each of the four treatment conditions. The number of CaMKIIα cells expressed in each section was normalized by the average number of CaMKIIα cells of the Near layer of both of the aCSF + dark and the CaMKIIα-PHOTAC + dark conditions (red dotted lines: 90% Confidence Interval of the dispersion of CaMKIIα expression). D) Comparisons of the slopes describing the loss of CaMKIIα with tissue depth that indicates that in the CaMKIIα-PHOTAC + 390 nm condition there is a gradual reduction of CaMKIIα-expressing neurons as we move towards the Near section. ANOVA comparing the slope revealed an effect of Condition (F_3,18_ = 13.13, p =.0001). Bonferroni-corrected post-hoc tests revealed significant differences between the CaMKIIα-PHOTAC + 390 nm condition and the control conditions (all p’s < 0.005) and no differences amongst the control conditions (all p’s > 0.9).

**Figure 5.**
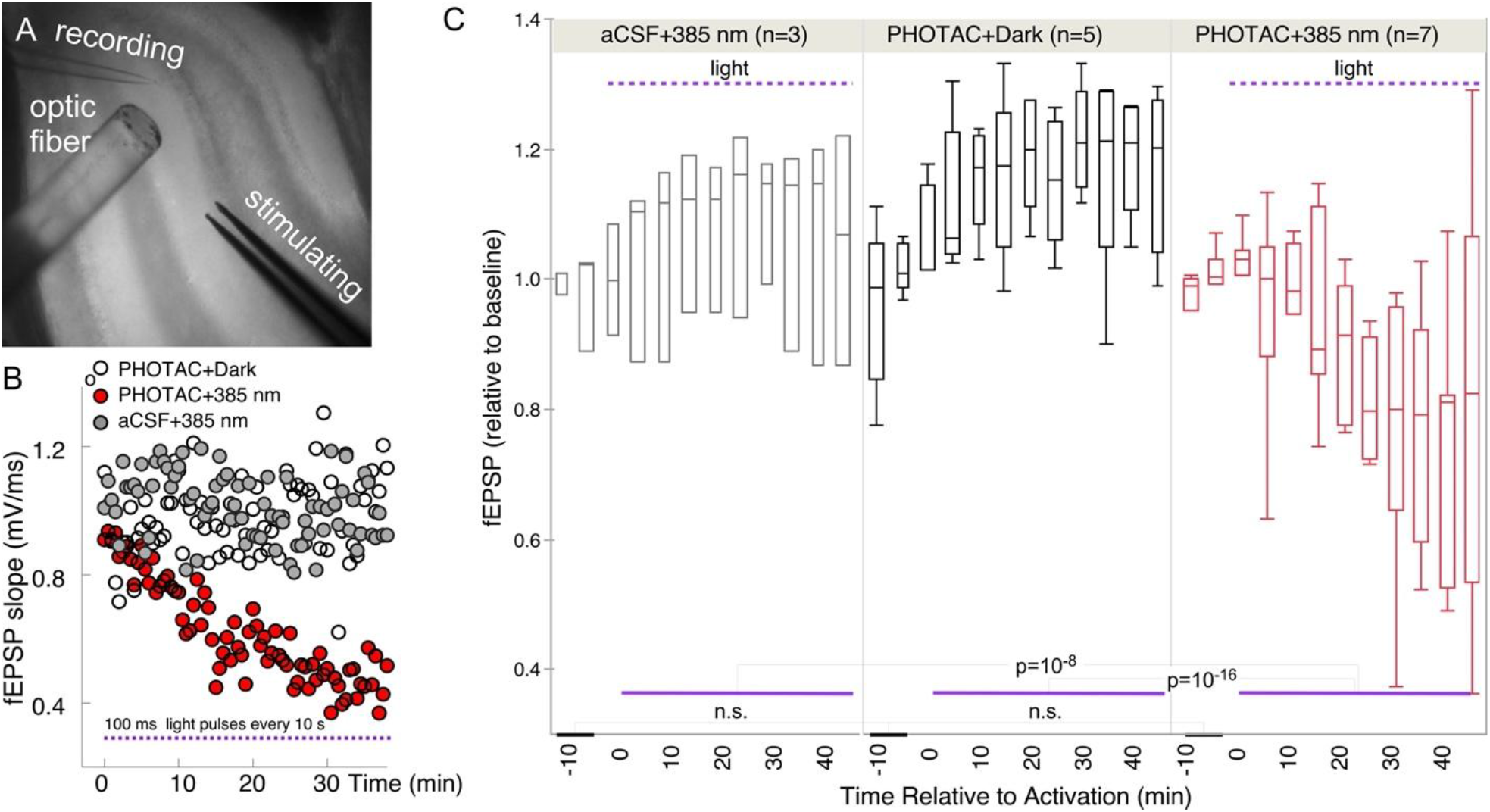
Optical Control of Synaptic Responses. A) Photograph illustrating the experimental configuration of the optic fibers and electrodes. B) Slopes of field excitatory postsynaptic potentials (fEPSP) in the presence and absence of **CaMKIIα-PHOTAC** and/or light. C) Statistical analysis of biological replicates.

First, we characterized the **CaMKIIα-PHOTAC** photophysical and thermal properties (Fig. 2). The compound isomerizes maximally to the biologically active *cis* configuration upon 385 and 390-nm illumination (Fig. 2B). Thermally, **CaMKIIα-PHOTAC** is relatively bistable, relaxing to its more stable *trans* form with a half-life of 8.86 hours at 37 °C in DMSO solution (Fig. 2C). Back-isomerization to the inactive form could be achieved by irradiation with wavelengths higher than 450 nm. In accordance with the logic of azobenzene photoswitches, numerous cycles of photochemical isomerization were possible without any observable photodegradation or fatigue (Fig. 2D).

Our design of a PHOTAC that targets proteins thought to be crucial for synaptic function is motivated by our interest in the molecular mechanisms of persistent memory storge CaMKIIα and PKM being persistent kinase candidates ^6^. Efforts to understand the role of these proteins have relied upon conventional pharmacology, pharmacogenetics, and deletion of their respective genes. But lack of pharmacological selectivity, as well genetic compensation and perinatal mortality have confounded those efforts ^13^, at times leading to incorrect conclusions ^39, 40^. As described here, PHOTAC pharmacology that can be activated and localized with light, and is not dependent on stoichiometric binding, may provide a powerful new class of tool for manipulations, although of course, the applications are not limited to neuroscience.

## Methods

### Chemical Synthesis

General information, experimental procedures, and characterization are summarized in the Supplementary Information.

### Protocols UV-Vis Spectroscopy

UV-Vis spectra were recorded on a Varian Cary 60 Scan UV-Vis spectrometer equipped with a Peltier PCB-1500 Thermostat and an 18-cell holder using Brand disposable UV cuvettes (70-850 μL, 10 mm light path) by Brandtech Scientific Inc. Sample preparation and all experiments were performed under red light conditions in a dark room. All UV-Vis measurements were performed with dimethyl sulfoxide (DMSO) as the solvent.

#### Wavelength scan

Light at different wavelengths was provided by an Optoscan Monochromator with an Optosource (75 mW lamp), which was controlled through a program written in Matlab. Irradiation to establish the photostationary state took place from the top through a fiber-optic cable. For each compound a 10 mM stock solution in DMSO was prepared and diluted to a 50 μM concentration prior to the experiment. Spectra with illumination were acquired from 550 to 370 nm in 20 nm steps going from higher to lower wavelengths and illuminating 5 minutes for each wavelength.

#### Thermal relaxation

Compounds were pre-irradiated with 390 nm light and observing the absorption at 370 nm over 12 hours at 37 °C in tightly sealed cuvettes.

### LED Illumination

#### Cell disco

For illumination of the slices during Western Blot and immunohistochemistry experiments, the cell disco system as previously described in the literature was used ^41^ with 390 nm light-emitting diodes (LEDs) purchased from Roithner Lasertechnik (VL390-5-15). Pulsed irradiation was performed using 100-ms pulses every 10 s controlled by an Arduino system.

### Tissue Preparation and PHOTAC Treatment

Acute mouse hippocampal brain slices: Eleven 6-9 month old wild type mice of either sex were deeply anesthetized with isofluorane (5% in 100% oxygen) for 2min, then quickly decapitated/ Their brain was removed and 300 μm horizontal hippocampal slices were prepared using a vibrating microtome (Leica VT 1000S) in a carbogenated (5% CO2, 95% O2) ice-cold, dissecting aCSF solution (mM: 125 NaCl, 2.5 KCl, 6.8 MgSO4, 0.5 CaCl2, 25 Glucose, 1.25 NaH2PO4, 25 NaHCO3, 1.25 NaH2PO4, 2 Na Pyruvate (319 mOsm)). Under environmental red-light, slices were then transferred to the incubation chambers with room-temperature, carbogenated recovery buffer (mM: 125 NaCl, 2.5 KCl, 1 MgSO4, 2 CaCl2, 25 Glucose, 1.25 NaH2PO4, 25 NaHCO3, 1.25 NaH2PO4, 2 Na Pyruvate (319 mOsm)), and incubated for 6 hours. For PHOTAC treatment groups, slices were incubated in a recovery buffer containing 3 μM of the PHOTAC compound. During the incubation, slices were placed in light-proof boxes and exposed to the lighting conditions specified in the experiment. Data from 9 mice were used.

### Immunoblotting Analysis

Under environmental red-light, treated slices were recovered and immediately homogenized into microtubes (Bel-Art™ ProCulture™ micro-tube homogenizer system) containing 200 μL of radioimmunoprecipitation assay buffer containing protease and phosphatase inhibitors. Protein concentration of the lysates was determined using BCA (Thermo Fisher Scientific). Samples were resolved under denaturing and reducing conditions using 4 to 12% bis-tris gels (NuPAGE) and transferred to a polyvinylidene fluoride membrane (Immobilon-P, Millipore). Membranes were blocked with 5% nonfat dried milk and incubated with CAMKIIα (1:1000, Invitrogen PA5-84083) and VINC (1:1000, Bethyl Laboratories A302-535A) primary antibodies overnight at 4 °C. After washing the membranes, secondary antibodies coupled with horseradish peroxidase were applied (Amersham-GE). Immunoreactive bands were visualized by enhanced chemiluminescence reagent (Thermo Fisher Scientific), and signal was acquired using the ChemiDoc Imaging System (Bio-Rad). ImageJ software was used for densitometric analysis of immunoblots. Data was analyzed using Prism version 9.11 (GraphPad Software Inc.).

### Immunohistochemistry

Under environmental red-light, incubated slices were immediately fixed in 4% PFA for 24 h. Subsequently, tissue sections were washed and permeabilized three times for 10 minutes with PBS containing 0.1% Tween20 (PBS-T) and blocked for 1.5 h at room temperature with 10% normal goat serum and PBS-T. Sections were then incubated overnight at 4 °C with the primary antibody rabbit anti-CaMKIIα polyclonal antibody (1:1000, Invitrogen PA5-84083), mouse anti-parvalbumin (1:1000, Millipore MAB1572) or mouse anti-MAP2 (1:500, Abcam Ab11268). After washing 3 times for 10 minutes each in PBS-T, the sections were incubated with goat anti-rabbit IgG (H+L) Cross-Adsorbed secondary antibody, Alexa Fluor Plus 488 (1:500, Invitrogen, A32731) and Alexa Fluor 594 goat anti-mouse IgG (H+L) (1:500, Molecular Probes, A11032) for 2 h at room temperature. After washing four times for 10 minutes in PBS, the sections were mounted on glass slides with 4’,6-diamidino-2-306 phenylindole (DAPI) Fluoromount-G (Southern Biotech) or Vectashield (Vector Laboratories) and coverslipped. The slices were then kept in the dark at 4 °C and later investigated using an upright Leica SP8 confocal microscope and analyzed using ImageJ (version 1.53a). For each slice, 8.5 μm-thick Z-stacks of the dorsal suprapyramidal and infrapyramidal blades of the dentate gyrus were created using the maximum intensity projection function in ImageJ, and measurements were made from each mouse in each region of interest.

### Slice Electrophysiology

Under environmental red-light, incubated slices were transferred to a submerged recording chamber in a Scientifica Slicescope apparatus and perfused with the same carbogenated aCSF solution as the recovery buffer at a rate of 5 ml/min and at 35-36 °C. Field excitatory postsynaptic potential (fEPSP) responses to a 100 μs, ∼200 mA stimulus from a bipolar electrode (FHC) in the supra-pyramidal blade of the dentate gyrus was recorded with a borosilicate glass pipette taper-pulled to an impedance of 5-7 MOhm (Sutter Instruments) and filled with the ACSF solution. The test pulse intensity was the intensity eliciting 70% of the maximal response given an input-output curve of seven intensities ranging from 0 μA to 300 μA. The slope of the synaptic response was measured using Clampfit 11.0.3. and normalized to each slice’s average baseline response.

#### Optic fiber

For illumination of the slices during synaptic recording, a 400 μm-diameter optic fiber was used. 385 mn light source was purchased from Thorlabs. Pulsed irradiation was performed using 100-ms pulses every 10 s controlled by a Master-8 A.M.P.I system.

## Author Contributions

TK, CJ, AGP, MR performed research and analyzed data; DT, AAF conceived of research, and wrote the manuscript with contributions from all authors.

