## Supplemental Information for "A Photoactivated Protein Degrader for Optical Control of Synaptic Function"

for

**Table of Contents**


| | |
|---|---|
| Supplemental Figures | S3 |
| General Information | S6 |
| Synthetic Procedures and Characterization | S7 |
| NMR spectra | S15 |
| References | S21 |

### Supporting Figures

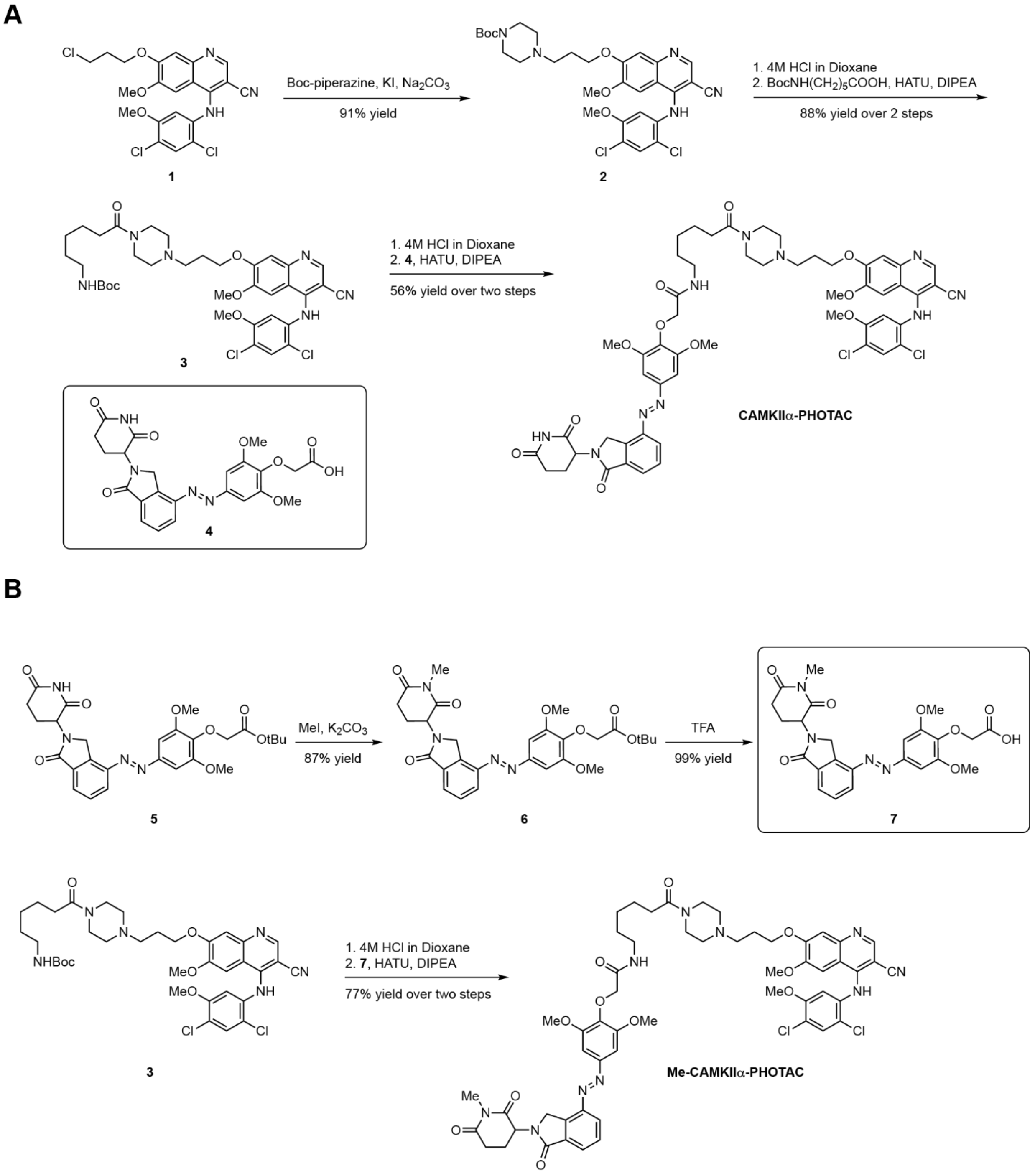


**Supporting Information Figure S1**. A) Synthesis of **CaMKIIα-PHOTAC**. B) Synthesis of **Me-CaMKIIα-PHOTAC.**

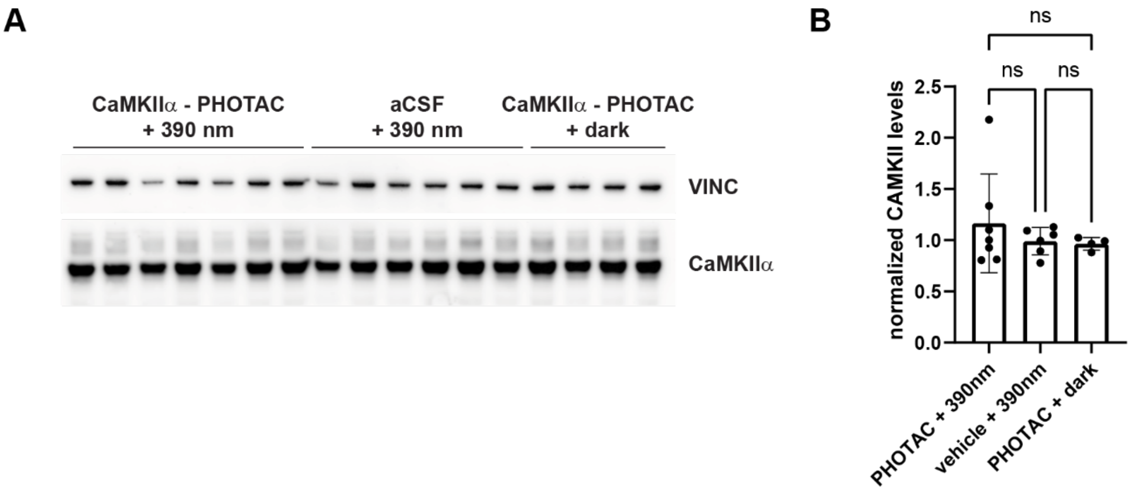


**Supporting Information Figure S2.** Western Blot analysis of CaMKII levels in hippocampal mice brain slices. A) Immunoblot of brain slices treated with either **CaMKIIa-PHOTAC** (3 μM) or aCSF (vehicle) under irradiation (390 nm, 100 ms every 10 s) or dark. B) Quantification of CaMKIIα levels normalized to VINC loading control.

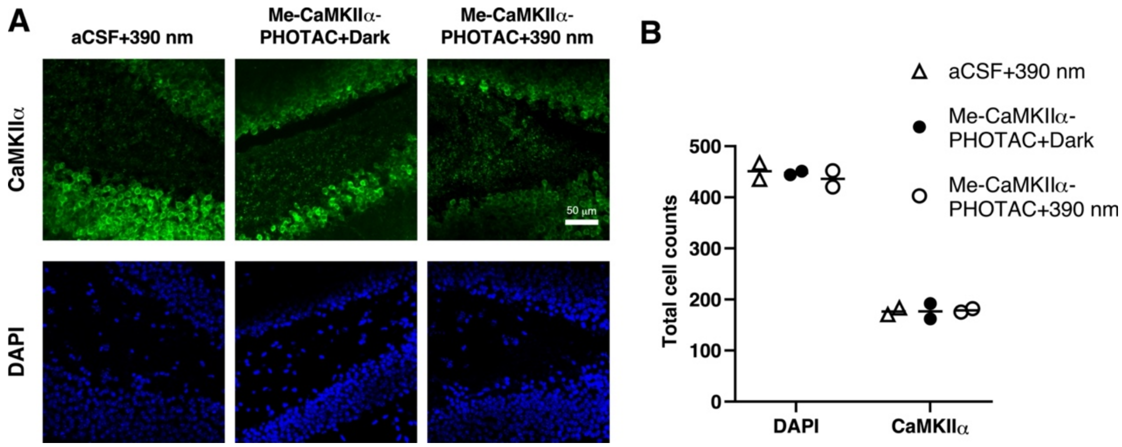


**Supporting Information Figure S3**. Immunohistochemistry of CaMKIIa and DAPI staining after control **Me-CaMKIIα-PHOTAC** treatment and activation. A) Examples of microphotographs of control slices incubated with either CaMKIIα-PHOTAC (3 μM) irradiated (390 nm, 100 ms every 10 s) or in the dark, or aCSF and irradiated (390 nm, 100 ms every 10 s). The tissue was later fixed and immunostained. B) Examples and quantitation of DAPI staining and immunohistochemistry for CaMKIIα.

### General Information

#### General experimental conditions

All reactions were conducted with magnetic stirring at room temperature unless otherwise noted. Reactions at elevated temperatures were performed using an oil bath or aluminum block as the heat transfer medium, with the reported temperature corresponding to that of the heat transfer medium.

#### Chemicals

All chemicals were obtained from common vendors and used without further purification unless otherwise noted. All solvents were purchased with "certified ACS" or higher quality. Anhydrous solvents were prepared with a solvent purification system by filtration of HPLC grade solvents through alumina or purchased from Acros Organics in AcroSeal™ bottles.

#### Chromatography

Retardation Factors (Rf) were determined by analytical thin-layer chromatography (TLC) performed on pre-coated glass plates from Millipore Sigma (TLC Silica Gel 60 Plates, 250 µm layer thickness, F254 fluorescence indicator), with visualization by exposure to ultraviolet light (254 nm) or by staining with a basic potassium permanganate solution. Column chromatography was performed with silica gel obtained from Millipore Sigma (Geduran® Si 60, 40 – 63 µm). Column chromatography was either performed manually or with an automated chromatography system (Teledyne Isco CombiFlash®).

#### Nuclear magnetic resonance (NMR) spectroscopy

Proton and carbon ($^{1}$H- and $^{13}$C-) NMR spectra was recorded on Bruker Avance III HD (400/100 MHz, with BBFO Cryoprobe™) spectrometer. All spectra were recorded at a temperature of 25 °C in 5 mm tubes in deuterated solvents purchased from Cambridge Isotope Laboratories, Inc. (chloroform-*d* or $CDCl_3$, 99.8% D; dimethyl sulfoxide-$d_6$ or DMSO-$d_6$, 99.9% D). For 1H-NMR spectra chemical shifts (δ) in parts per million (ppm) relative to tetramethylsilane (δ = 0 ppm) are reported using the residual protic solvent ($CHCl_3$ in $CDCl_3$: δ = 7.26 ppm, DMSO-$d_5$ in DMSO-$d_6$: δ = 2.50 ppm) as an internal reference. For $^{13}$C-NMR spectra, chemical shifts in ppm relative to tetramethylsilane (δ = 0 ppm) are reported using the central resonance of the solvent signal ($CDCl_3$: δ = 77.16 ppm, DMSO-$d_6$: δ = 39.52 ppm) as an internal reference. The abbreviations used for multiplicities and descriptors are s = singlet, d = doublet, t = triplet, q = quartet or combinations thereof, m = multiplet and br = broad. NMR spectral data was analyzed with the program MestreNova.

#### Mass spectrometry (MS)

**Liquid Chromatography-Mass Spectometry (LCMS):** Samples were measured on an Agilent Technologies 1260 II Infinity connected to an Agilent Technologies 6120 Quadrupole mass spectrometer with ESI ionization source. Elution was performed using a gradient from 5:95% to 100:0% MeCN:$H_2O$ with 0.1% formic acid over 5 min, if not indicated otherwise.

**High-resolution mass spectra (HRMS):** Spectra were obtained with an Agilent 6224 Accurate Mass time-of-flight (TOF) LC/MS system using an electrospray ionization (ESI) ion source. All reported data refers to positive ionization mode.

**Synthetic procedures and characterization**

tert-butyl 4-(3-((3-cyano-4-((2,4-dichloro-5-methoxyphenyl)amino)-6-methoxyquinolin-7-yl)oxy)propyl)piperazine-1-carboxylate (**2**)

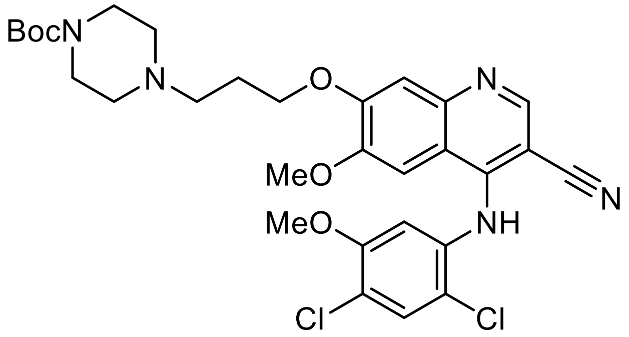


7-(3-chloropropoxy)-4-((2,4-dichloro-5-methoxyphenyl)amino)-6-methoxyquinoline-3-carbonitrile (**1**, 163.0 mg, 0.349 mmol, 1.0 eq.), Boc-piperazine (97.6 mg, 0.524 mmol, 1.5 eq.), potassium iodide (115.9 mg, 0.698 mmol, 2 eq.), and sodium carbonate (222.1 mg, 2.095 mmol, 6.0 eq.) were dissolved in *n*-butanol (3.5 ml). The solution was stirred and heated to 105°C for 24 hours. The solution was cooled to room temperature and concentrated under reduced pressure. The resulting residue was dissolved in ethyl acetate (10 ml), washed with water (10 ml) and saturated aqueous sodium chloride (10 ml). The organic layer was dried over sodium sulfate and concentrated under reduced pressure. The residue was purified by column chromatography (0 to 7% methanol in dichloromethane) to afford the title compound (195.6 mg, 0.317 mmol, 91%) as beige solid.

**$R_f$** = 0.57 (7.5% methanol in dichloromethane)

**$^1$H-NMR** (400 MHz, $CDCl_3$) δ 8.71 (s, 1H), 7.49 (s, 1H), 7.44 (s, 1H), 6.90 (s, 1H), 6.72 (s, 1H), 6.46 (s, 1H), 4.27 (t, J = 6.6 Hz, 2H), 3.78 (s, 3H), 3.67 (s, 3H), 3.50 – 3.35 (m, 4H), 2.65 – 2.35 (m, 6H), 2.19 – 2.04 (m, 2H), 1.46 (s, 9H).

**$^{13}$C-NMR** (100 MHz, $CDCl_3$) δ 154.80, 154.45, 154.01, 150.44, 149.91, 147.87, 147.63, 137.15, 130.75, 118.49, 117.18, 116.53, 115.05, 110.09, 105.43, 101.28, 94.63, 79.96, 67.58, 56.72, 56.22, 55.02, 53.10, 28.57.

| | | |
|---|---|---|
| **LCMS** (ESI): | $t_{ret}$ = 2.840 min | 616.2 m/z $[M+H]^+$. |
| **HRMS** (ESI): | calc. for $C_{30}H_{35}Cl_2N_5O_5Na^+$: | 638.1907 m/z $[M+Na]^+$. |
| | found: | 638.1879 m/z $[M+Na]^+$. |

tert-butyl (6-(4-(3-((3-cyano-4-((2,4-dichloro-5-methoxyphenyl)amino)-6-methoxyquinolin-7-yl)oxy)propyl)piperazin-1-yl)-6-oxohexyl)carbamate (**3**)

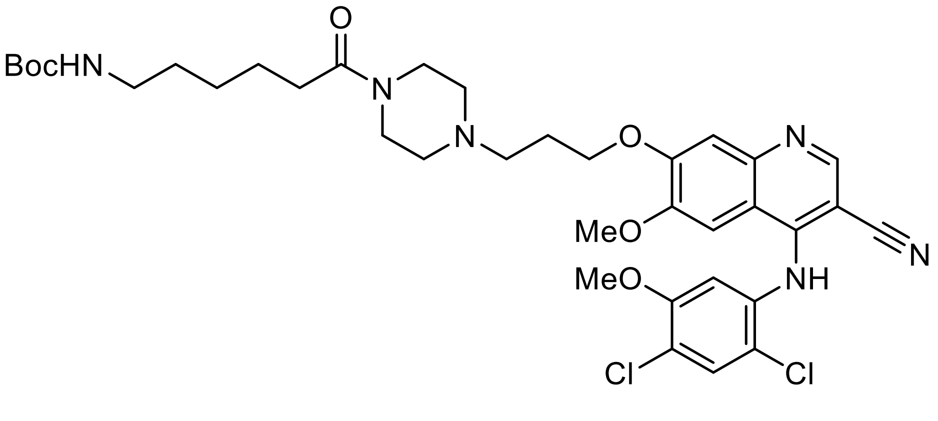


tert-butyl 4-(3-((3-cyano-4-((2,4-dichloro-5-methoxyphenyl)amino)-6-methoxyquinolin-7-yl)oxy)propyl)piperazine-1-carboxylate (**2**, 100.5 mg, 0.163 mmol, 1.0 eq.) was dissolved in DCM (3.4 ml) until all the substrate was dissolved. 4 M HCl in dioxane was added dropwise (1.0 ml) to the solution. Solids immediately began precipitating and the solution was stirred for an hour. The mixture was concentrated under reduced pressure and to the resulting solid, 6-((tert-butoxycarbonyl)amino) hexanoic acid (45.2 mg, 0.195 mmol, 1.2 eq.), HATU (92.8 mg, 0.244 mmol, 1.5 eq.) were added under nitrogen atmosphere. The solids were then dissolved in anhydrous DMF (11.6 ml), the mixture was stirred until all solids dissolved and distilled DIPEA (11.6 µl, 0.651 mmol, 4.0 eq) was added dropwise. The solution was stirred overnight at room temperature and concentrated under reduced pressure until most of the DMF had evaporated. The resulting residue was dissolved in 10 ml of ethyl acetate and 10 ml of 10% LiCl aqueous solution were added. Subsequently, the layers were separated, and the aqueous layer was further extracted (3 x 10 ml) with ethyl acetate. The organic layers were combined, washed with saturated aqueous sodium chloride (10 mL) and dried over sodium sulfate. The solvent was removed under reduced pressure and the residue purified by column chromatography (0 to 10% methanol in dichloromethane) to afford the title compound (104.1 mg, 0.143 mmol, 88%) as a beige solid.

**$R_f$** = 0.51 (7.5% methanol in dichloromethane)

**$^1$H-NMR** (400 MHz, $CDCl_3$) δ 8.70 (s, 1H), 7.49 (s, 1H), 7.43 (s, 1H), 6.92 (s, 1H), 6.78 (s, 1H), 6.48 (s, 1H), 4.56 (s, 1H), 4.28 (t, J = 6.5 Hz, 2H), 3.78 (s, 3H), 3.67 (s, 3H), 3.65 – 3.47 (m, 4H), 3.11 (q, J = 6.4 Hz, 2H), 2.68 – 2.43 (m, 6H), 2.32 (t, J = 7.5 Hz, 2H), 2.18 – 2.09 (m, 2H), 1.68 – 1.60 (m, 2H), 1.54 – 1.46 (m, 2H), 1.43 (s, 9H), 1.40 – 1.32 (m, 2H).

**$^{13}$C-NMR** (100 MHz, $CDCl_3$) δ 171.43, 156.17, 154.46, 153.84, 150.36, 149.91, 147.83, 137.03, 130.74, 118.77, 117.59, 116.48, 115.01, 109.98, 105.90, 101.38, 94.24, 79.23, 67.29, 56.74, 56.25, 54.95, 53.40, 52.82, 40.50, 33.15, 30.04, 28.58, 26.68, 24.94.

| | | |
|---|---|---|
| **LCMS** (ESI): | $t_{ret}$ = 3.061 min | 729.3 m/z $[M+H]^+$. |
| **HRMS** (ESI): | calc. for $C_{36}H_{46}Cl_2N_6O_6Na^+$: | 751.2787 m/z $[M+Na]^+$. |
| | found: | 751.2748 m/z $[M+Na]^+$. |

**CAMKIIα-PHOTAC**

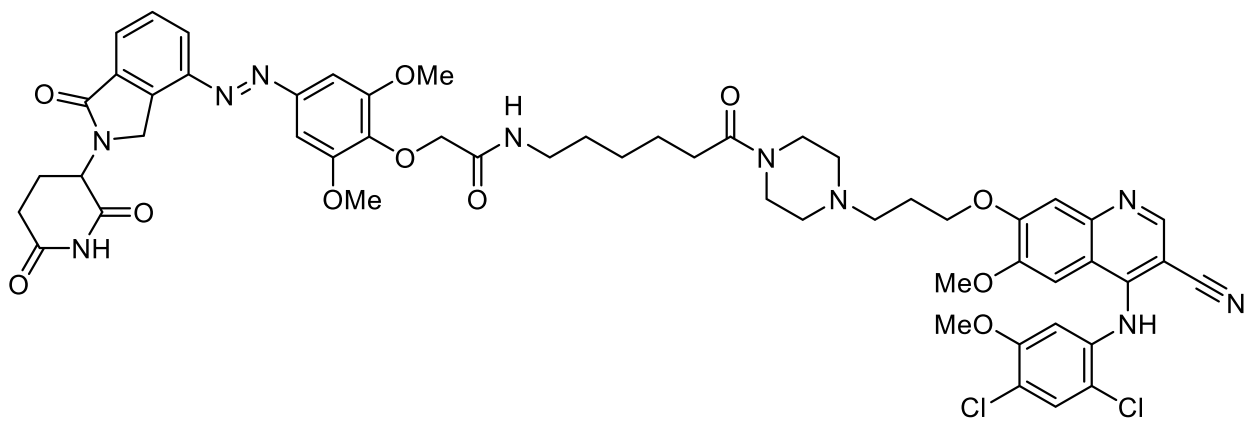

tert-butyl (6-(4-(3-((3-cyano-4-((2,4-dichloro-5-methoxyphenyl)amino)-6-methoxyquinolin -7-yl)oxy)propyl)piperazin-1-yl)-6-oxohexyl)carbamate (**3**, 16.8 mg, 0.023 mmol, 1.0 eq.) was dissolved in DCM (0.5 ml) until all the substrate was dissolved. 4 M HCl in dioxane was added dropwise (0.15 ml) to the solution. Solids immediately began precipitating and the solution was stirred for an hour. The mixture was concentrated under reduced pressure and to the resulting solid, (*E*)-2-(4-((2-(2,6-dioxopiperidin-3-yl)-1-oxoisoindolin-4-yl)diazenyl)-2,6-dimethoxyphenoxy) acetic acid [1] (**4**, 16.3 mg, 0.034 mmol, 1.5 eq.), HATU (12.8 mg, 0.034 mmol, 1.5 eq.) were added under nitrogen atmosphere. The solids were then dissolved in anhydrous DMF (1.6 ml), the mixture was stirred until all solids dissolved and distilled DIPEA (1.6 µl, 0.90 mmol, 4.0 eq) was added dropwise. The solution was stirred overnight at room temperature and concentrated under reduced pressure until most of the DMF had evaporated. The resulting residue was dissolved in 2 ml of a 10:1 DCM and methanol solution and 4 ml of 10% LiCl aqueous solution were added. Subsequently, the layers were separated, and the aqueous layer was further extracted (3 x 2 ml) with the DCMand methanol solution. The organic layers were combined, washed with saturated aqueous sodium chloride (4 mL) and dried over sodium sulfate. The solvent was removed under reduced pressure and the residue purified by column chromatography (0 to 10% methanol in dichloromethane) to afford the title compound (13.8 mg, 0.013 mmol, 56%) as an orange solid.

$\mathbf{R_f}$ = 0.6 (7.5% methanol in dichloromethane)

**$^1$H-NMR** (400 MHz, DMSO-*d*$_6$) δ 11.04 (s, 1H), 9.63 (s, 1H), 8.41 (s, 1H), 8.21 (dd, J = 7.8, 1.0 Hz, 1H), 7.94 (s, 1H), 7.91 (d, J = 1.1 Hz, 1H), 7.84 (s, 1H), 7.79 (t, J = 7.7 Hz, 1H), 7.74 (s, 1H), 7.36 (s, 2H), 7.33 (s, 2H), 5.16 (dd, J = 13.2, 5.1 Hz, 1H), 4.85 – 4.65 (m, 2H), 4.44 (s, 2H), 4.20 (t, J = 6.4 Hz, 2H), 3.96 – 3.93 (m, 9H), 3.86 (s, 3H), 3.49 – 3.35 (m, 4H), 3.18 (q, J = 6.6 Hz, 2H), 2.99 – 2.89 (m, 1H), 2.67 – 2.59 (m, 1H), 2.57 – 2.52 (m, 1H), 2.48 – 2.33 (m, 6H), 2.28 (t, J = 7.4 Hz, 2H), 2.10 – 2.02 (m, 1H), 2.00 – 1.94 (m, 2H), 1.54 – 1.44 (m, 4H), 1.32 – 1.25 (m, 2H).

**$^{13}$C-NMR** (100 MHz, DMSO) δ 174.48, 172.90, 171.01, 170.42, 167.92, 167.14, 153.99, 152.72, 152.57, 150.80, 149.38, 148.27, 146.34, 145.75, 139.26, 136.19, 134.50, 133.82, 129.83, 129.63, 128.64, 125.48, 123.06, 120.35, 116.93, 113.51, 112.45, 109.38, 101.84, 100.60, 86.33, 71.86, 66.74, 56.80, 56.36, 56.25, 54.12, 53.01, 52.48, 51.85, 48.30, 44.89, 40.93, 38.10, 32.15, 31.28, 28.96, 26.10, 25.74, 24.50, 22.32.

**LCMS** (ESI): $t_{ret}$ = 3.110 min 1093.4 m/z $[M+H]^+$.

| | | |
|---|---|---|
| **HRMS** (ESI): | calc. for $C_{54}H_{58}Cl_2N_{10}O_{11}Na^+$: | 1115.3556 m/z $[M+Na]^+$. |
| | found: | 1115.3539 m/z $[M+Na]^+$. |

tert-butyl (E)-2-(2,6-dimethoxy-4-((2-(1-methyl-2,6-dioxopiperidin-3-yl)-1-oxoisoindolin-4-yl)diazenyl)phenoxy)acetate (**6**)

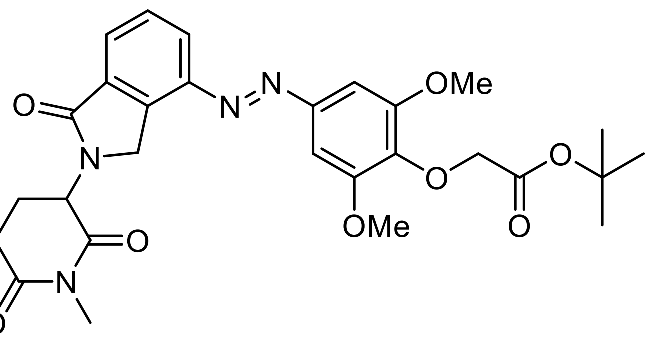


Tert-butyl (E)-2-(4-((2-(2,6-dioxopiperidin-3-yl)-1-oxoisoindolin-4-yl)diazenyl)-2,6-dime thoxyphenoxy)acetate (**5**, 200.0 mg, 0.371 mmol, 1.0 eq.) was dissolved in DMF (2.0 ml) and THF (5.1 ml) under a nitrogen atmosphere. To the solution, methyl iodide (79.1 mg, 0.557 mmol, 1.5 eq.) and potassium carbonate (154.0 mg, 1.114 mmol, 3.0 eq.) were added. The solution was stirred and heated to 65°C for 24 hours. The solution was cooled to room temperature and concentrated under reduced pressure. The resulting residue was dissolved in ethyl acetate (50 ml) and water (50 ml). The layers were separated and the aqueous layer was extracted (2 x 50 ml) and saturated aqueous sodium chloride (50 ml). The organic layer was dried over sodium sulfate and concentrated under reduced pressure. The residue was purified by column chromatography (0 to 5% methanol in dichloromethane) to afford the title compound (178.5 mg, 0.323 mmol, 87%) as orange solid.

$\mathbf{R_f}$ = 0.65 (5% methanol in dichloromethane)

**$^1$H-NMR** (400 MHz, $CDCl_3$) δ 8.19 – 8.16 (m, 1H), 8.01 – 7.97 (m, 1H), 7.72 – 7.67 (m, 1H), 7.19 (s, 2H), 5.22 (dd, J = 13.7, 5.1 Hz, 1H), 4.81 – 4.69 (m, 2H), 4.67 (s, 2H), 3.96 (s, 6H), 3.21 (s, 2H), 3.06 – 2.83 (m, 2H), 2.51 – 2.18 (m, 2H), 1.48 (s, 9H).

**$^{13}$C-NMR** (100 MHz, $CDCl_3$) δ 171.25, 170.11, 168.57, 168.20, 152.73, 148.30, 146.91, 139.83, 134.11, 133.46, 129.44, 129.39, 125.96, 100.76, 81.84, 81.75, 81.73, 69.93, 69.86, 56.49, 56.45, 52.68, 48.30, 32.14, 28.12, 27.25, 22.82.

**LCMS** (ESI): $t_{ret}$ = 4.101 min 553.2 m/z $[M+H]^+$.

**HRMS** (ESI): calc. for $C_{28}H_{32}N_4O_8Na^+$: 575.2112 m/z $[M+Na]^+$.
found: 575.2129 m/z $[M+Na]^+$.

(E)-2-(2,6-dimethoxy-4-((2-(1-methyl-2,6-dioxopiperidin-3-yl)-1-oxoisoindolin-4-yl)diazenyl)phenoxy)acetic acid (**7**)

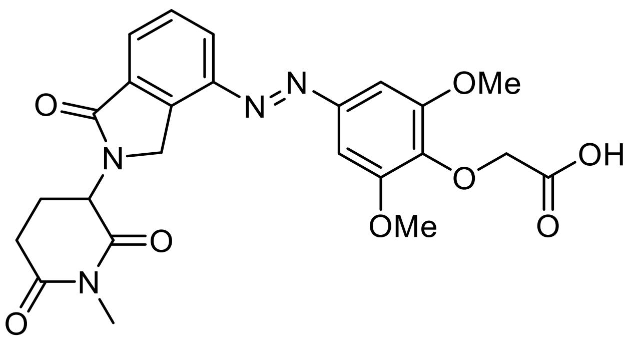


tert-butyl (E)-2-(2,6-dimethoxy-4-((2-(1-methyl-2,6-dioxopiperidin-3-yl)-1-oxoisoindolin-4-yl)diazenyl)phenoxy)acetate (**6**, 101.0 mg, 0.183 mmol, 1.0 eq.) was dissolved in DCM (1.0 ml) and TFA (1.0 ml) was added dropwise. The solution turned from orange to red upon addition of TFA. After 1 hour, the reaction was concentrated under reduced pressure. The solution was further dried under high vacuum for 48 hours to obtain the title compound (90.0 mg, 0.181 mmol, 99%) as orange solid.

$\mathbf{R_f}$ = 0.14 (7.5% methanol in dichloromethane)

**$^1$H-NMR** (400 MHz, $CDCl_3$) δ 8.22 – 8.17 (m, 1H), 8.05 – 8.00 (m, 1H), 7.72 (t, J = 7.7 Hz, 1H), 7.23 (s, 2H), 5.24 (dd, J = 13.6, 5.0 Hz, 1H), 4.85 – 4.70 (m, 2H), 4.70 (s, 2H), 4.02 (s, 6H), 3.21 (s, 3H), 3.08 – 2.82 (m, 2H), 2.50 – 2.17 (m, 2H).

**$^{13}$C-NMR** (100 MHz, $CDCl_3$) δ 171.23, 170.12, 170.11, 168.54, 152.36, 149.38, 146.71, 139.03, 139.00, 134.18, 133.51, 129.80, 129.52, 126.51, 100.39, 71.08, 56.57, 52.72, 48.34, 32.12, 27.27, 22.83.

| | | |
|---|---|---|
| **LCMS** (ESI): | $t_{ret}$ = 3.067 min | 497.1 m/z $[M+H]^+$. |
| **HRMS** (ESI): | calc. for $C_{24}H_{25}N_4O_8^+$: | 497.1667 m/z $[M+H]^+$. |
| | found: | 497.1703 m/z $[M+H]^+$. |

**Me-CAMKIIα-PHOTAC**

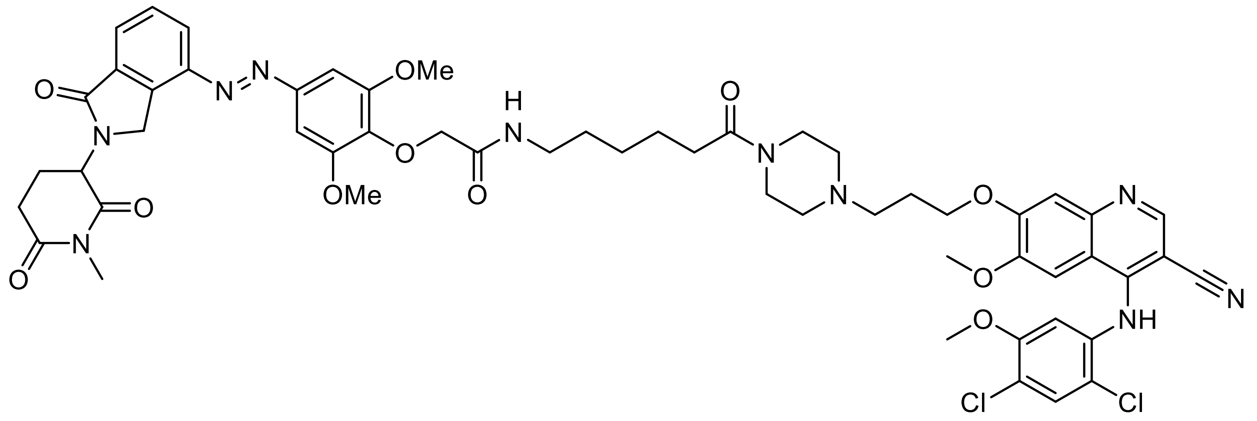

tert-butyl (6-(4-(3-((3-cyano-4-((2,4-dichloro-5-methoxyphenyl)amino)-6-methoxyquinolin -7-yl)oxy)propyl)piperazin-1-yl)-6-oxohexyl)carbamate (**3**, 16.8 mg, 0.023 mmol, 1.0 eq.) was dissolved in DCM (0.5 ml) until all the substrate was dissolved. 4 M HCl in dioxane was added dropwise (0.15 ml) to the solution. Solids immediately began precipitating and the solution was stirred for an hour. The mixture was concentrated under reduced pressure and to the resulting solid, (E)-2-(2,6-dimethoxy-4-((2-(1-methyl-2,6-dioxopipe ridin-3-yl)-1-oxoisoindolin-4-yl)diazenyl)phenoxy)acetic acid (16.8 mg, 0.034 mmol, 1.5 eq.), HATU (12.8 mg, 0.034 mmol, 1.5 eq.) were added under nitrogen atmosphere. The solids were then dissolved in anhydrous DMF (1.6 ml), the mixture was stirred until all solids dissolved and distilled DIPEA (1.6 µl, 0.90 mmol, 4.0 eq) was added dropwise. The solution was stirred overnight at room temperature and concentrated under reduced pressure until most of the DMF had evaporated. The resulting residue was dissolved in 2 ml of a 10:1 DCM and methanol solution and 4 ml of 10% LiCl aqueous solution were added. Subsequently, the layers were separated, and the aqueous layer was further extracted (3 x 2 ml) with the DMC and methanol solution. The organic layers were combined, washed with saturated aqueous sodium chloride (4 mL) and dried over sodium sulfate. The solvent was removed under reduced pressure and the residue purified by column chromatography (0 to 10% methanol in dichloromethane) to afford the title compound (19.2 mg, 0.017 mmol, 77%) as an orange solid.

**$R_f$** = 0.63 (7.5% methanol in dichloromethane)

**$^1$H-NMR** (400 MHz, DMSO-$d_6$) δ 9.60 (s, 1H), 8.41 (s, 1H), 8.21 (dt, J = 7.8, 1.5 Hz, 1H), 7.95 – 7.90 (m, 2H), 7.82 (s, 1H), 7.79 (t, J = 7.7 Hz, 1H), 7.74 (s, 1H), 7.36 (s, 2H), 7.33 (s, 2H), 5.22 (dd, J = 25.0, 13.4, 5.1 Hz, 1H), 4.87 – 4.65 (m, 2H), 4.43 (s, 2H), 4.19 (t, J = 6.4 Hz, 2H), 3.94 (s, 9H), 3.86 (s, 3H), 3.50 – 3.36 (m, 4H), 3.18 (q, J = 6.7 Hz, 2H), 3.01 (s, 3H), 3.00 – 2.96 (m, 1H), 2.83 – 2.75 (m, 1H), 2.62 – 2.53 (m, 1H), 2.48 – 2.32 (m, 6H), 2.28 (t, J = 7.4 Hz, 2H), 2.13 – 2.03 (m, 1H), 2.01 – 1.91 (m, 2H), 1.55 – 1.43 (m, 4H), 1.32 – 1.24 (m, 2H).

**$^{13}$C-NMR** (100 MHz, DMSO) δ 172.38, 171.11, 170.87, 168.38, 167.66, 154.47, 153.23, 153.04, 151.28, 149.85, 148.74, 146.85, 146.24, 139.75, 136.67, 135.14, 134.98, 134.29, 130.31, 130.12, 128.86, 126.00, 123.55, 120.84, 117.40, 113.99, 112.90, 109.85, 102.28, 101.10, 86.80, 72.33, 67.25, 57.28, 56.84, 56.70, 54.64, 53.55, 53.01, 52.84, 48.71, 45.35, 41.39, 38.58, 32.63, 31.89, 29.43, 27.10, 26.57, 26.28, 24.99, 22.06.

**LCMS** (ESI): $t_{ret}$ = 3.230 min 1107.4 m/z $[M+H]^+$.

| | | |
|---|---|---|
| **HRMS** (ESI): | calc. for $C_{55}H_{60}Cl_2N_{10}O_{11}Na^+$: | 1129.3712 m/z $[M+Na]^+$. |
| | found: | 1129.3695 m/z $[M+Na]^+$. |

$^{1}$H-NMR (400 MHz, $CDCl_3$) – **2**

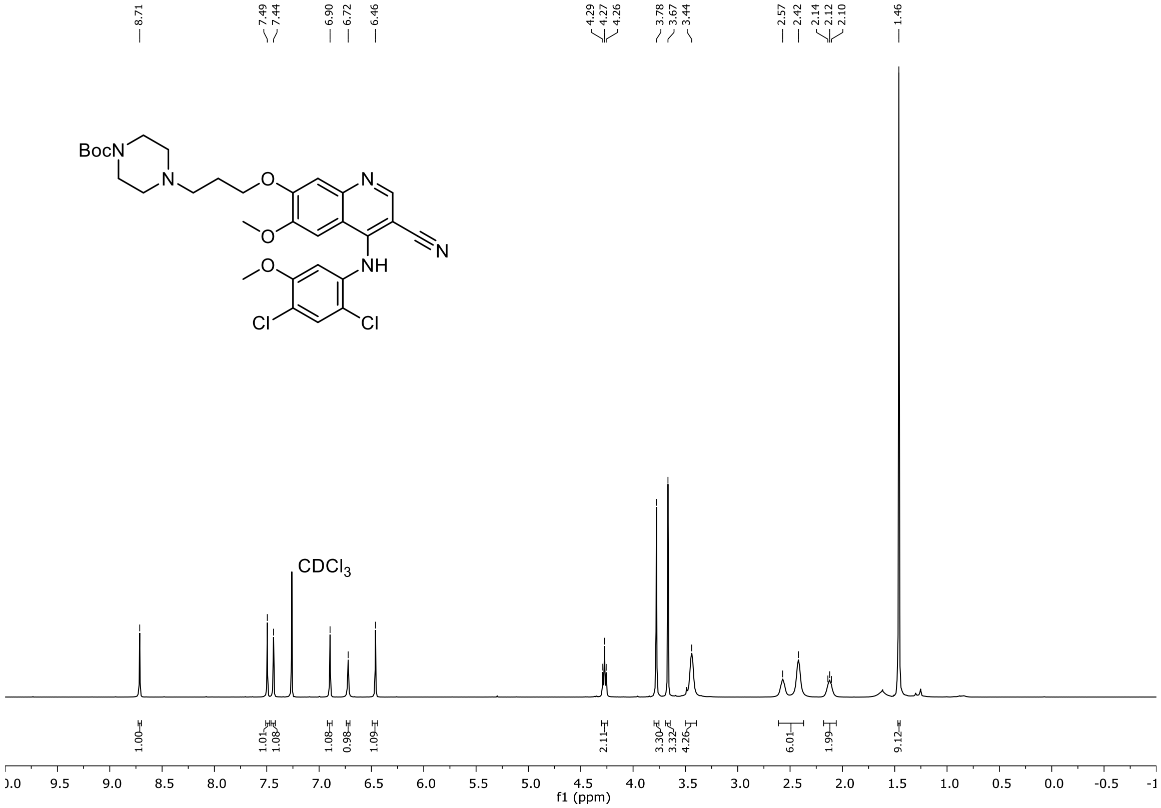


$^{13}$C-NMR (100 MHz, $CDCl_3$) – **2**

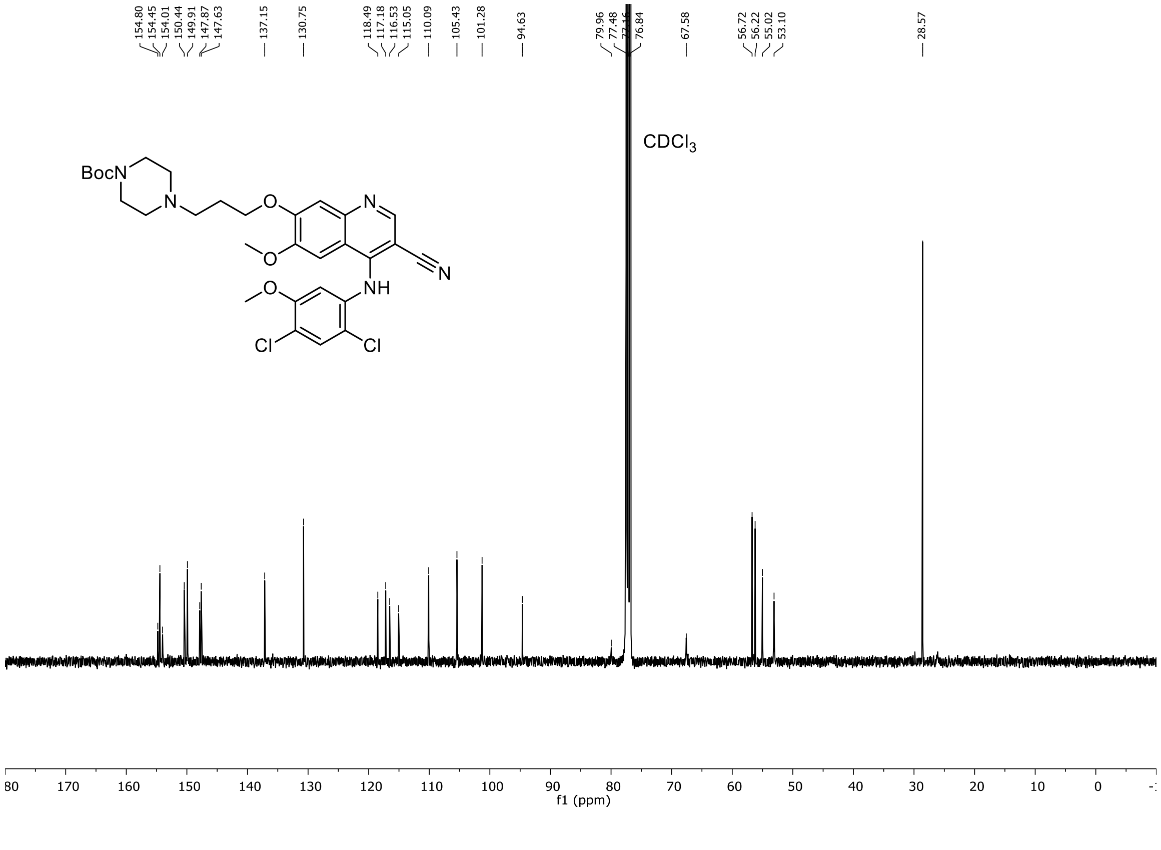

$^1$H-NMR (400 MHz, $CDCl_3$) – **3**

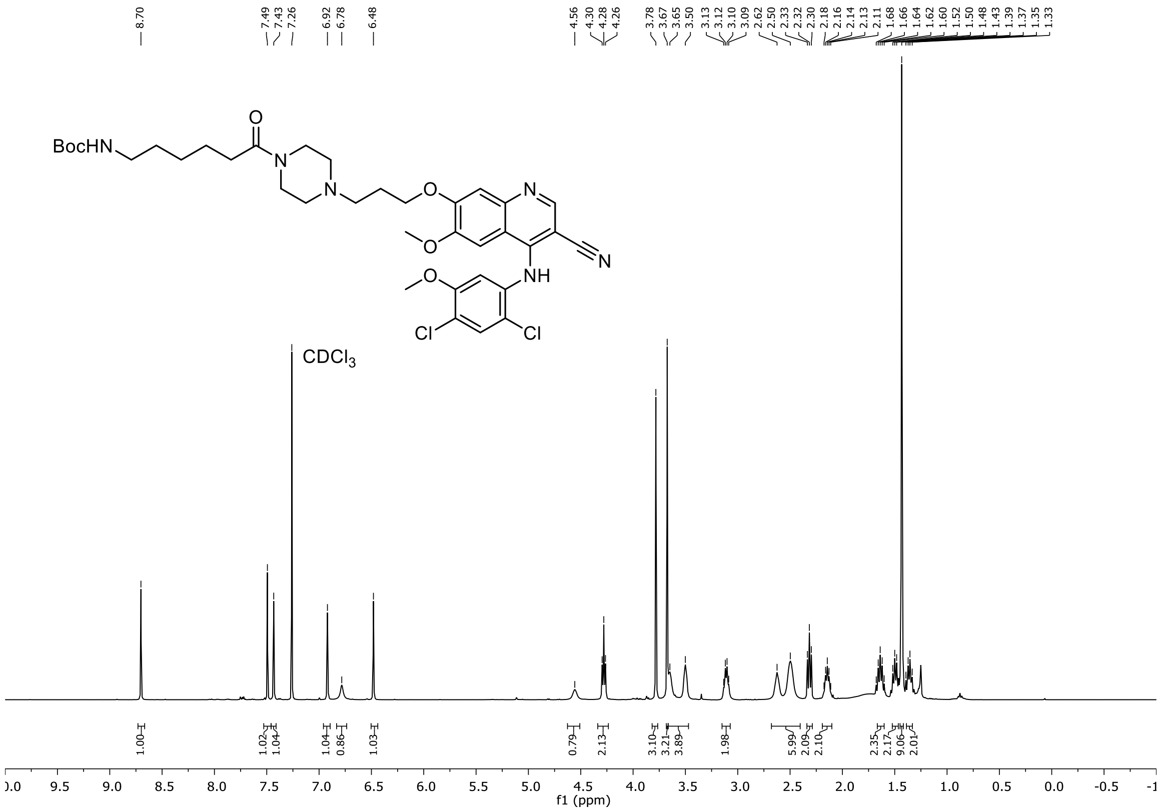

$^{13}$C-NMR (100 MHz, $CDCl_3$) – **3**

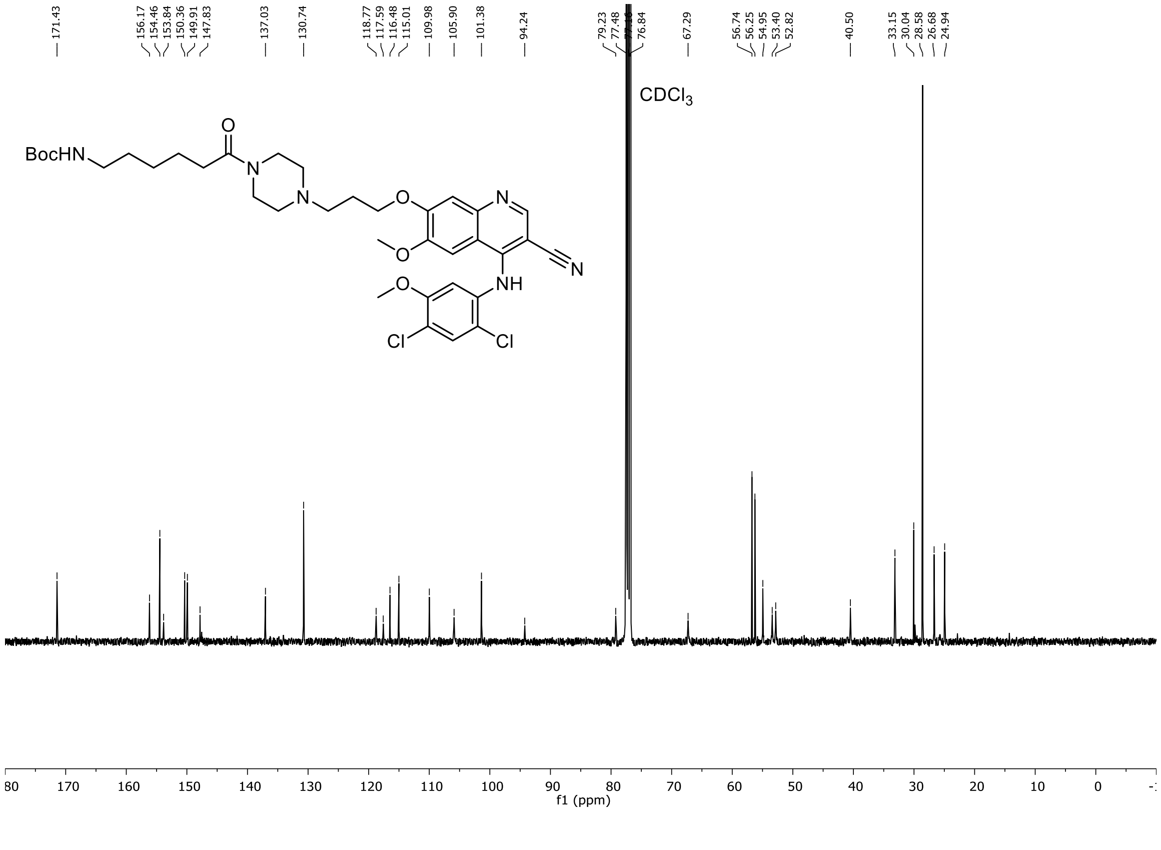

$^{1}$H-NMR (400 MHz, DMSO-$d_6$) – **CaMKIIα-PHOTAC**

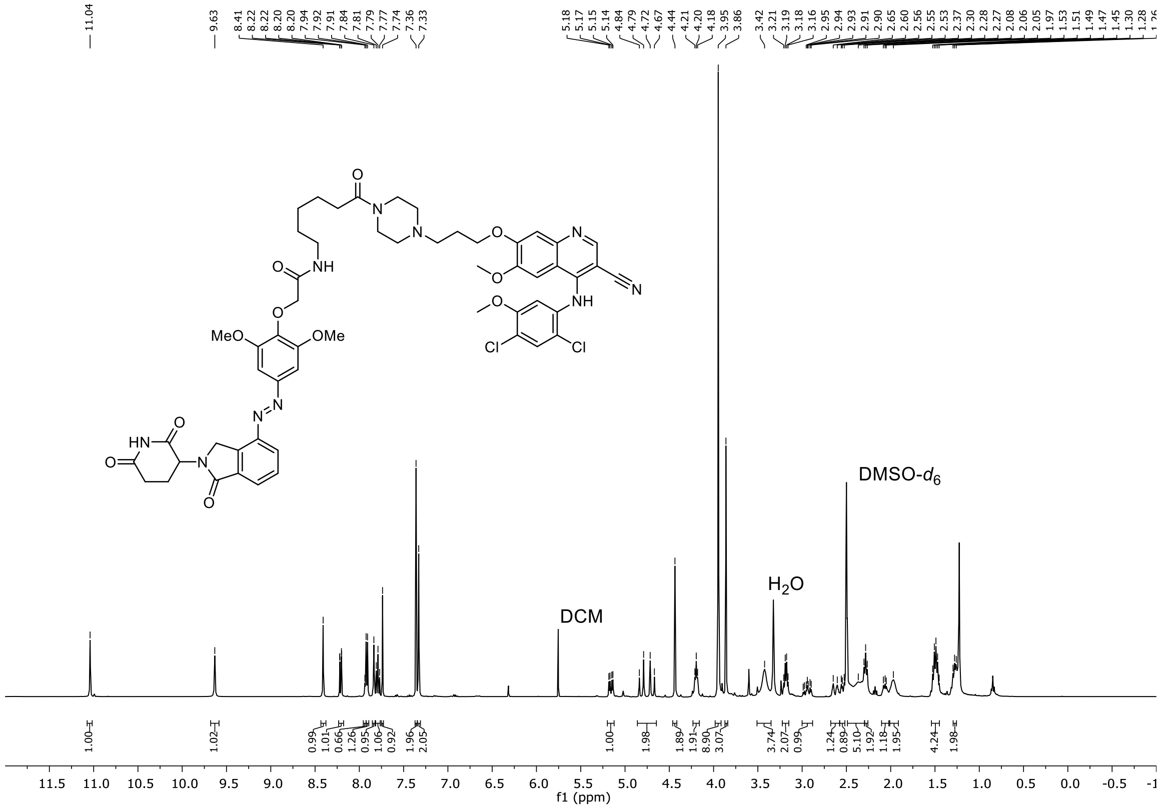


$^{13}$C-NMR (100 MHz, DMSO-$d_6$) – **CaMKIIα-PHOTAC**

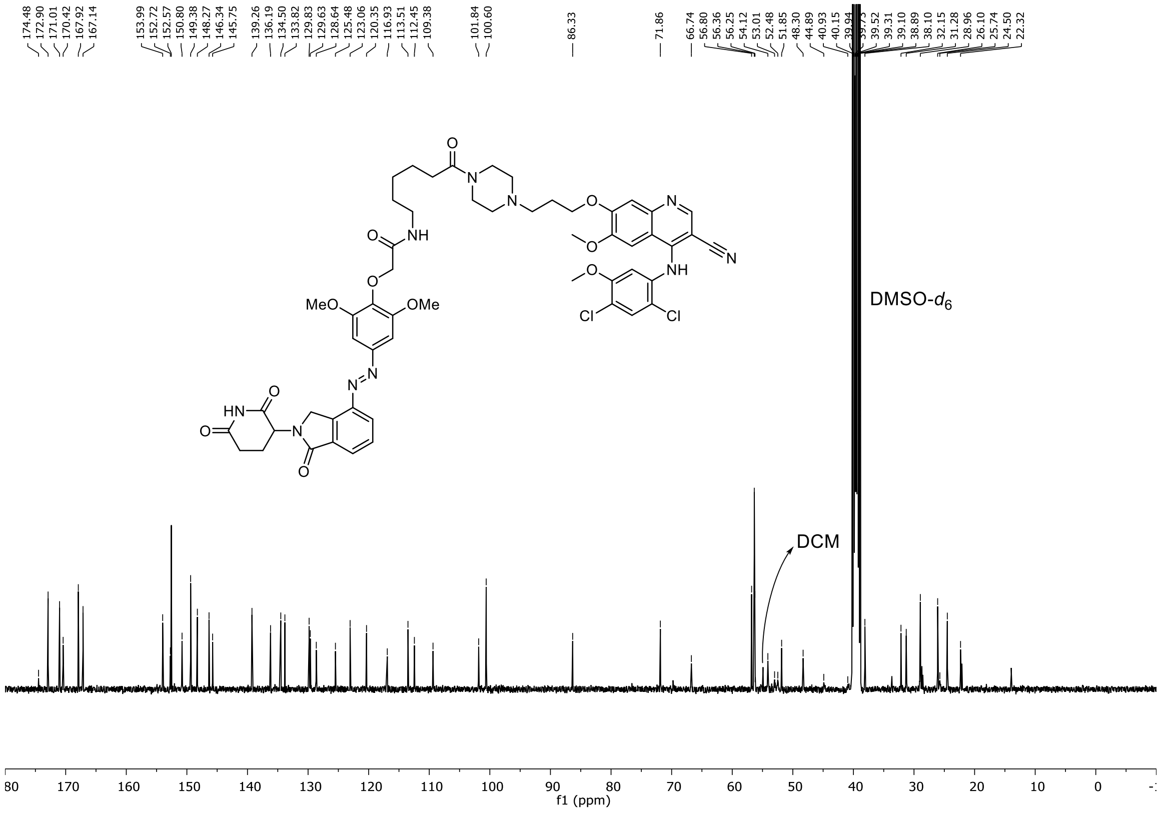

$^{1}$H-NMR (400 MHz, $CDCl_3$) – **6**

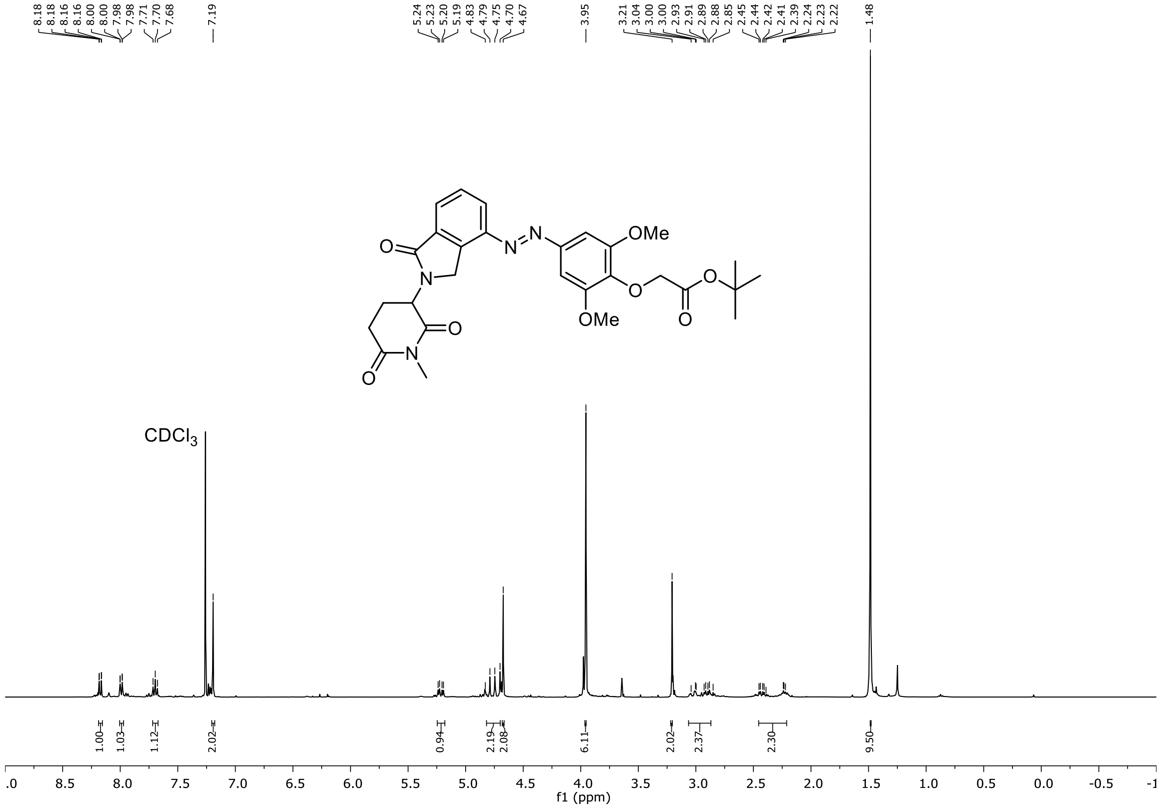


$^{13}$C-NMR (100 MHz, $CDCl_3$) – **6**

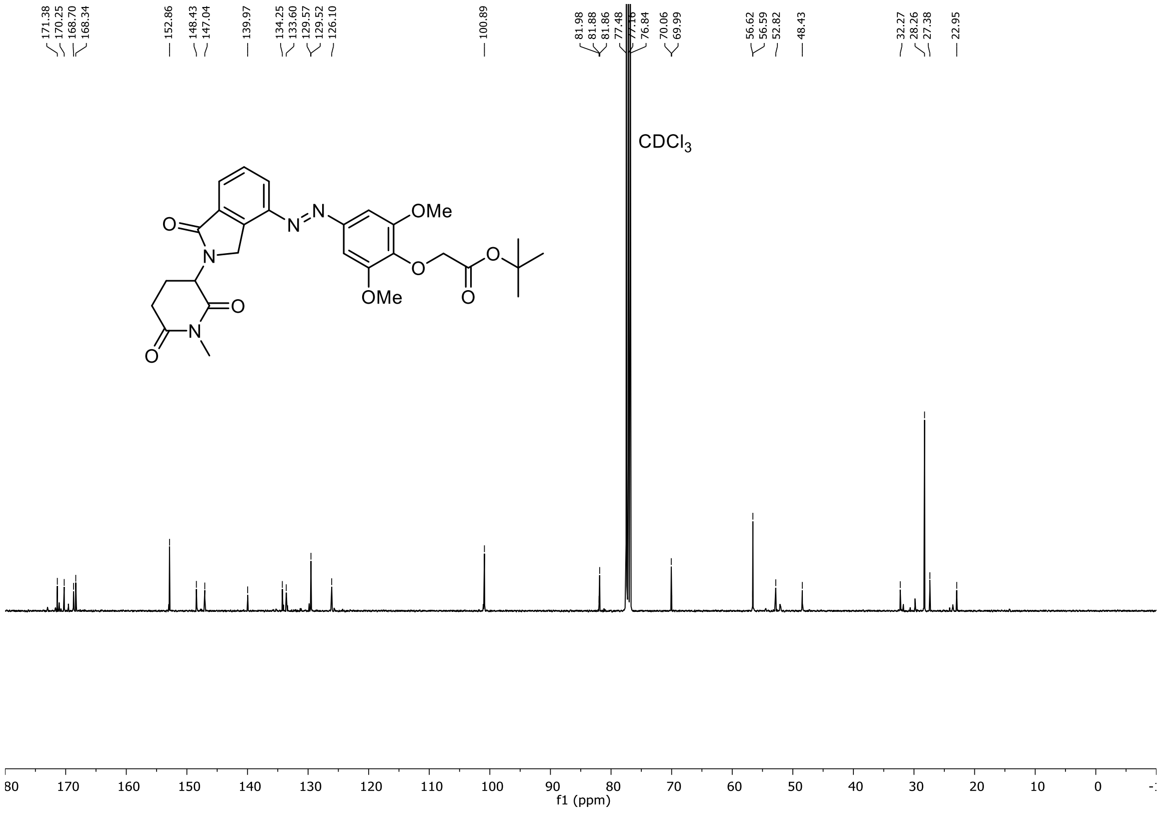

$^{1}$H-NMR (400 MHz, $CDCl_3$) – **7**

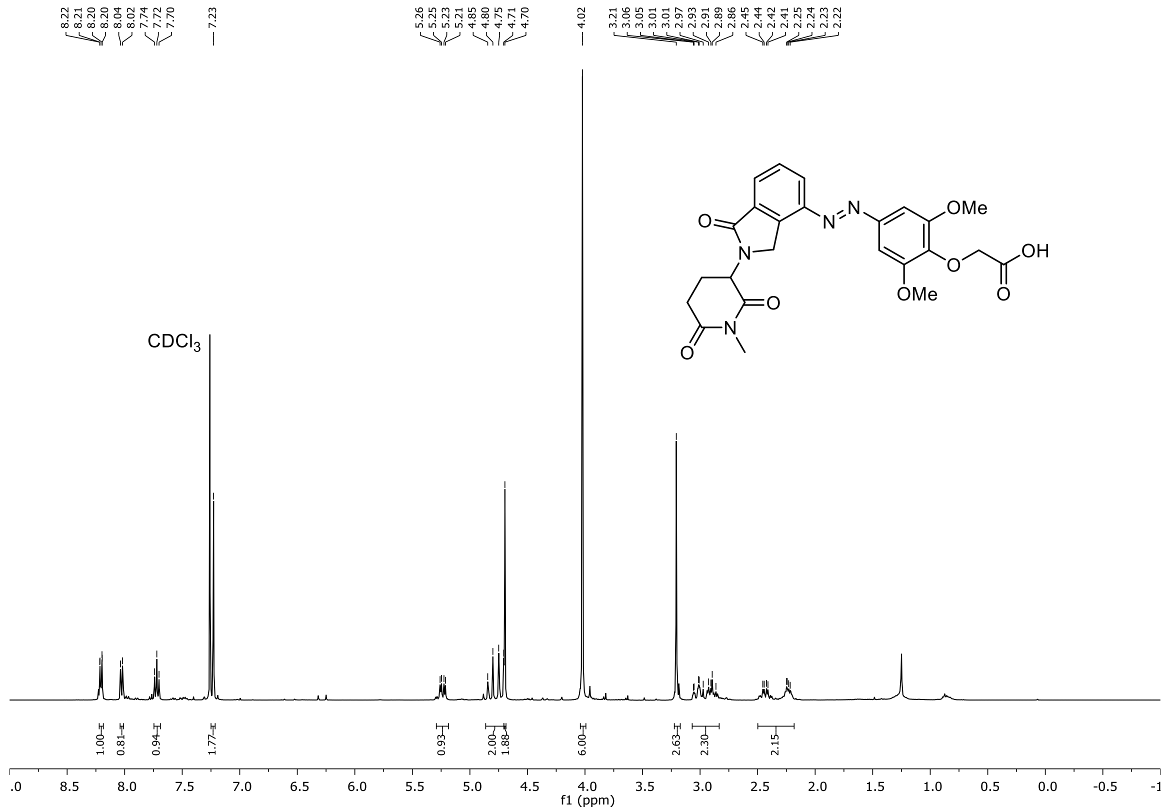

$^{13}$C-NMR (100 MHz, $CDCl_3$) – **7**

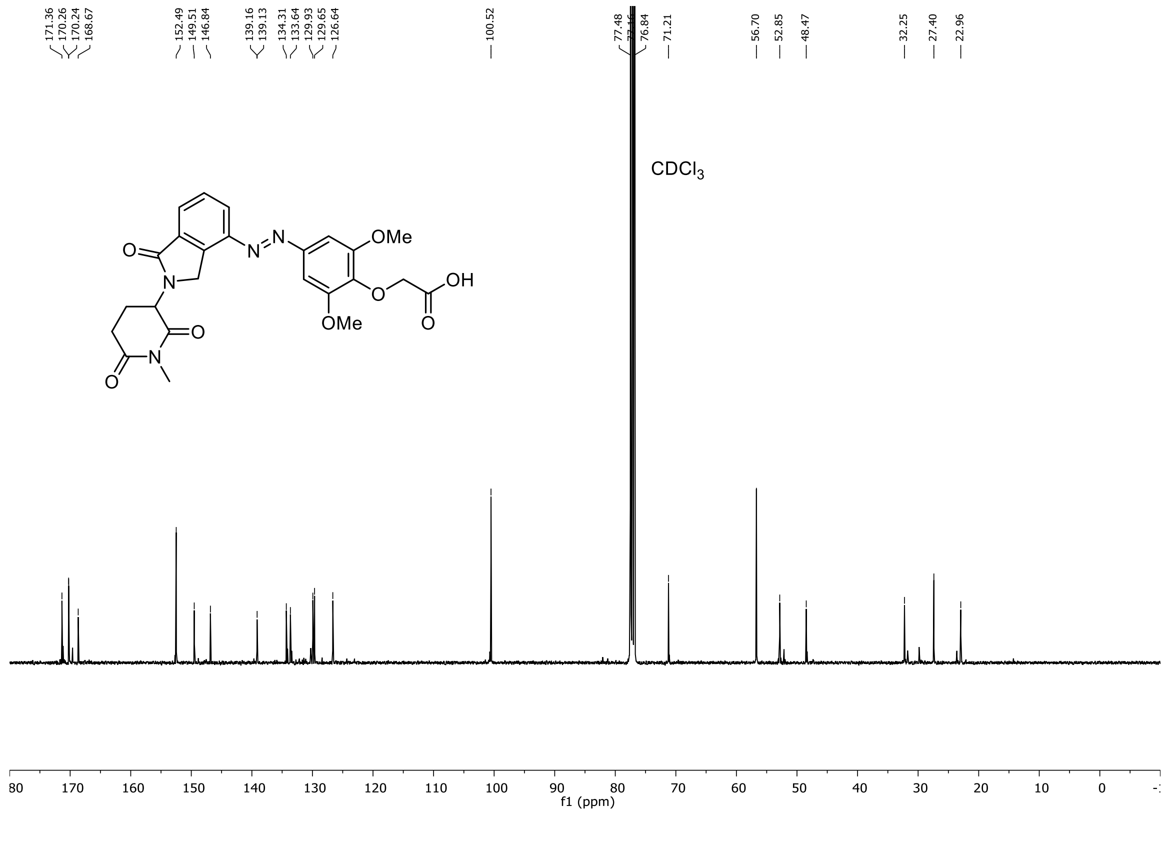

$^{1}$H-NMR (400 MHz, DMSO-$d_6$) – **Me-CaMKIIα-PHOTAC**

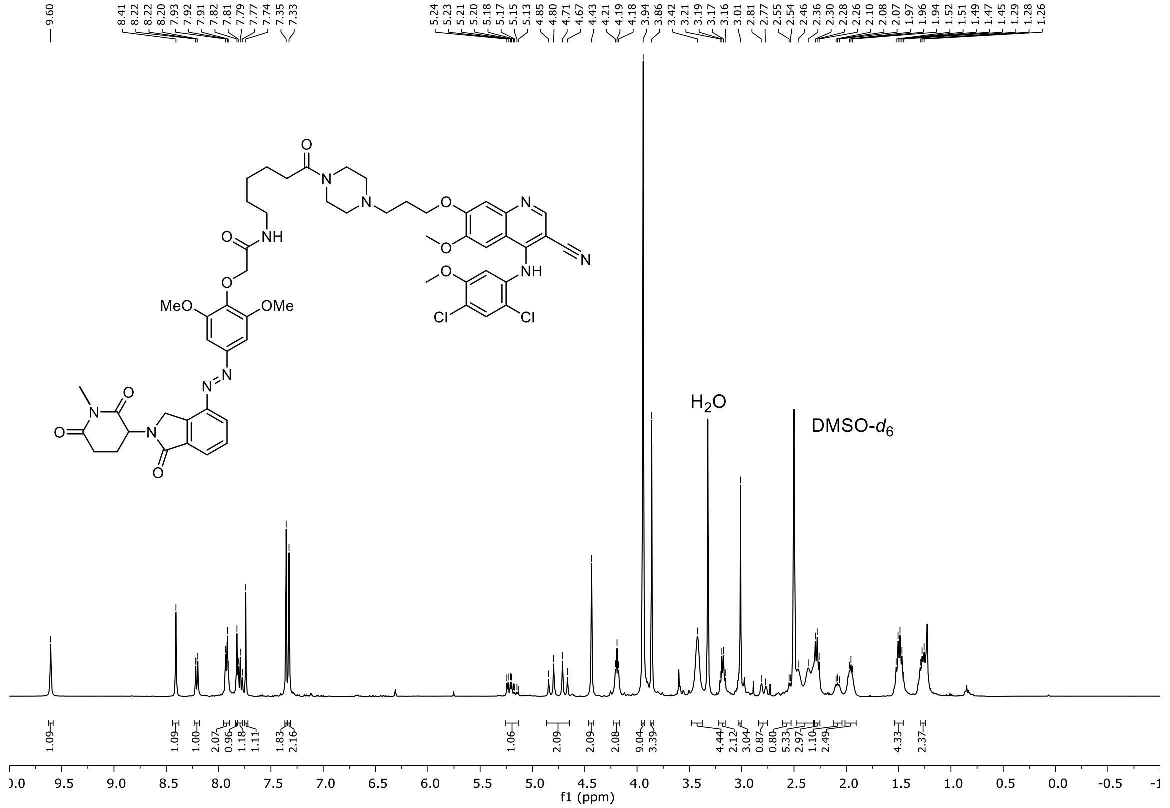

$^{13}$C-NMR (100 MHz, DMSO-$d_6$) – **Me-CaMKIIα-PHOTAC**

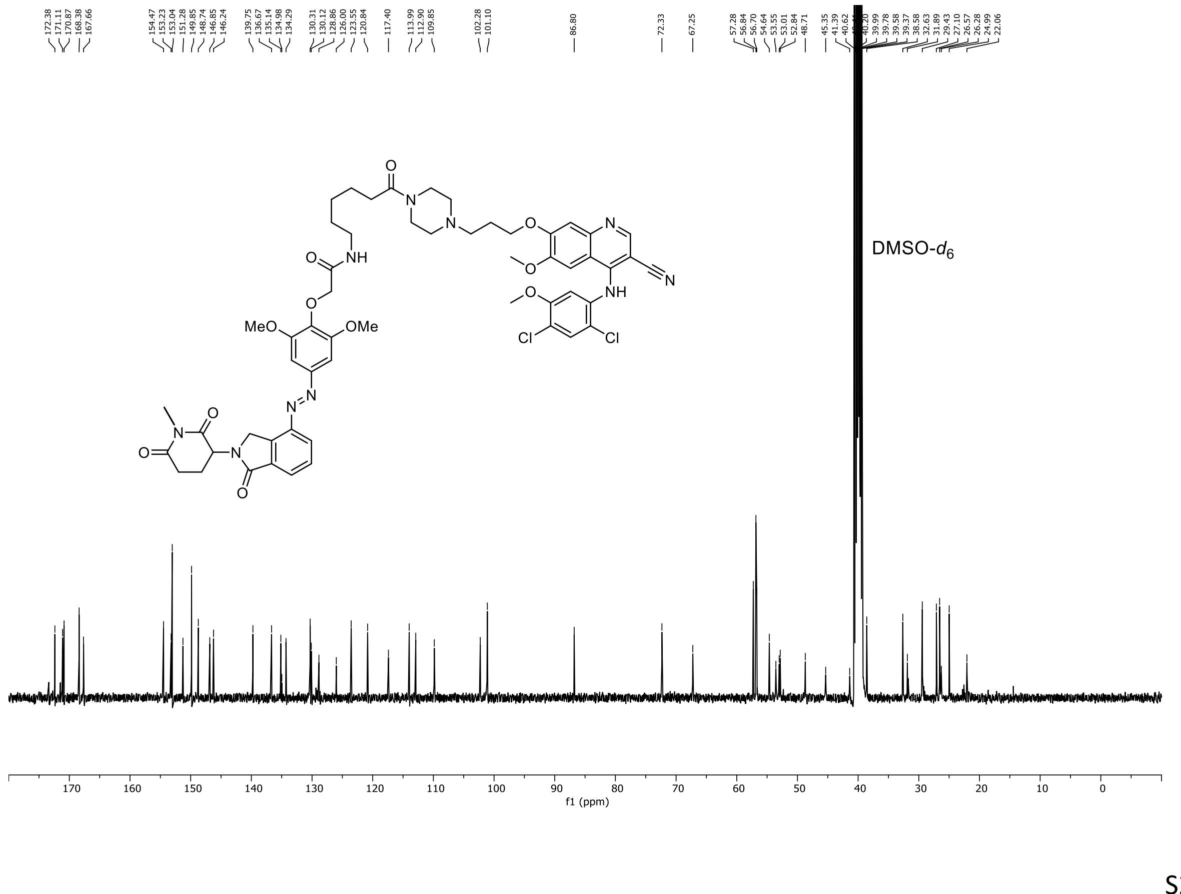
